# The Landscape of Receptor-Mediated Precision Cancer Combination Therapy: A Single-Cell Perspective

**DOI:** 10.1101/2020.01.28.923532

**Authors:** Saba Ahmadi, Pattara Sukprasert, Rahulsimham Vegesna, Sanju Sinha, Fiorella Schischlik, Natalie Artzi, Samir Khuller, Alejandro A. Schäffer, Eytan Ruppin

## Abstract

The availability of single-cell transcriptomics data opens new opportunities for rational design of combination cancer treatments. Mining such data, we employed combinatorial optimization techniques to explore the landscape of optimal combination therapies in solid tumors including brain, head and neck, melanoma, lung, breast and colon cancers. We assume that each individual therapy can target any one of 1269 genes encoding cell surface receptors, which may be targets of CAR-T, conjugated antibodies or coated nanoparticle therapies. As a baseline case, we studied the killing of at least 80% of the tumor cells while sparing more than 90% of the non-tumor cells in each patient, as a putative regimen. We find that in most cancer types, personalized combinations composed of at most four targets are then sufficient. However, the number of distinct targets that one would need to assemble to treat all patients in a cohort accordingly would be around 10 in most cases. Further requiring that the target genes be also lowly expressed in healthy tissues uncovers qualitatively similar trends. However, as one asks for more stringent and selective killing beyond the baseline regimen we focused on, we find that the number of targets needed rises rapidly. Emerging individual promising receptor targets include *PTPRZ1*, which is frequently found in the optimal combinations for brain and head and neck cancers, and *EGFR*, a recurring target in multiple tumor types. In sum, this systematic single-cell based characterization of the landscape of combinatorial receptor-mediated cancer treatments establishes first of their kind estimates on the number of targets needed, identifying promising ones for future development.

## Introduction

Personalized oncology offers hope that each patient’s cancer can be treated based on its genomic characteristics^1,2^. Several trials have suggested that it is possible to collect genomics data fast enough to inform treatment decisions^3–5^. Meta-analysis of Phase I clinical trials completed during 2011-2013 showed that overall, trials that used molecular biomarker information to influence treatment plans gave better results than trials that did not^6^. However, most precision oncology treatments utilize only one or two medicines, and resistant clones frequently emerge, emphasizing the need to deliver personalized medicine as multiple agents combined^6–11^. Important opportunities to combine systems biology and design of nanomaterials have been recognized to deliver medicines in combination to overcome drug resistance and combine biological effects^12^.

Here, we propose and study a new conceptual framework for designing future precision oncology treatments. It is motivated by the growing recognition that tumors typically have considerable intra-tumor heterogeneity (ITH)^13,14^ and thus need to be targeted with a combination of medicines such that as many as possible tumor cells are hit by at least one medicine. Our analysis is based on two recently emerging technologies: (1) the advancement of single-cell transcriptomics and proteomics measurements from patients’ tumors, which is anticipated to gradually enter into clinical use^15^, and (2) the introduction of “modular” treatments that target specific overexpressed genes/proteins to recognize cells in a specific manner and then use either the T cell immune response or a lethal toxin to kill the tumor cells preferentially.

Based on these two technological foundations, we formulate and systematically answer two basic translational questions. First, how many targeted treatments are needed to selectively kill most tumor cells while sparing most of the non-tumor cells in a given patient? And second, given a cohort of patients to treat, how many distinct single-target treatments need to be prepared beforehand to treat each patient effectively with the per-patient minimum number of targeted treatments?

We focus our analysis on genes encoding receptors on the cell surface, as these may be precisely targeted by any one of at least six technologies, including CAR-T therapy^16^, immunotoxins ligated to antibodies^17–18^, immunotoxins ligated to mimicking peptides^19^ and conventional chemotherapy ligated to nanoparticles^20^. These treatments are all termed “modular”, as they include one part that specifically targets the tumor cell via a gene/protein overexpressed on its surface and another part, the cytotoxic mechanism that kills the cells.

Two recent genome-wide analyses of modular therapies have focused on CAR-T therapy^21,22^, so we focus first on this technology to put our work in context. In the original formulation, CAR-T therapy used one cell surface target that marks the cells of interest, such as CD19 as a marker for B cells. To date, CAR-T therapy has been effective in achieving remissions for some blood cancers^16,23^, but less effective for solid tumors. MacKay et al.^22^ focused primarily on single targets and looked at combinations of two targets and did all analysis *in silico*. Dannenfelser et al.^23^ focused on predicting combinations of two and three targets and did most of their work *in silico*, with *in vitro* validation of two high-scoring predicted combinations in renal cancer. Importantly, these studies have analyzed *bulk tumor and normal expression* data to identify likely targets. Here we present the first analysis that aims to identify modular targets based on the analysis of *tumor single-cell transcriptomics*. This enables to study the research questions at a higher resolution but presents new analytical challenges that need to be addressed.

Two related difficulties with CAR-T therapy are i) toxicity to non-cancer cells^24,25^ and ii) difficulty in finding single targets that are sufficiently selective^22^. To address the toxicity problem, MacKay et al.^22^ selected 533 targets that had low expression in most tissues in the Genotype-Tissue Expression (GTEx) data; however, their analysis did not require that the targets are cell surface proteins. We proceed in a stepwise manner; we start with a formal analysis of a space of 1269 candidate cell surface receptors. Then, we add a low-expression requirement like that of MacKay et al.^22^. For completeness, we also tested their set of 533 genes.

To address the selectivity problem, various groups have engineered composite forms of CAR-T treatments that implement Boolean AND, OR, and NOT gates that have been tested for combinations of up to three target proteins^26–30^. Both MacKay et al. and Dannenfelser et al. presented *in silico* methods focusing on AND gates and pairs or trios of targets; Dannenfelser et al. analyzed 2538 likely cell surface proteins that are not necessarily receptors. We have chosen to focus on the simpler logical OR construction because that can be achieved not only by CAR-T technology^27,28^, but can also be implemented via other modular receptor-mediated treatment technologies by combining multiple single-target treatments, assuming that the composite treatment kills a cell if any one of the single treatments kill the cell. Conceptually, such a logical OR combination treatment can still achieve selectivity by choosing targets, each of which is expressed on a much higher proportion of cancer cells than non-cancer cells. One of our key contributions is to show that by using techniques from combinatorial optimization, one can find such effective combinations involving many targets, while previous studies were limited to at most three targets.

Beyond CAR-T, our analysis applies to several additional types of modular treatment technologies that rely instead on receptor-mediated endocytosis (RME) delivering a toxin via a targeted receptor to enter the cell^31,32^. These RME-based technologies include, e.g., conjugated antibodies and toxin delivering nanoparticles. Like CAR-T, these technologies do not downregulate the target receptor. For RME technologies and other technologies that work intracellularly, we anticipate combining modular treatments from one technology such that *all treatments use the same toxin or mechanism of cell killing*, thereby mitigating the need to test for interaction effects between pairs of different treatments.

To address this research challenge, we designed and implemented a computational approach named MadHitter (after the Mad Hatter from *Alice in Wonderland*) to identify optimal precision combination treatments that target membrane receptors (**Figure 1, A-C**). We define three key parameters related to the stringency of killing the tumor and protecting the non-tumor cells and explore how the optimal treatments vary with those parameters (**Figure 1B, C**). Solving this problem is analogous to solving the classical “hitting set problem” in combinatorial algorithms^33^, which is formally defined in the **Methods** section (see also **Supplementary Materials 1**). Unlike the previous studies on CAR-T targets, we define the problem in a personalized manner, intending that each patient will get optimal treatments for her or his tumor from among a collection of treatments available at the cohort level.

**Figure 1.**
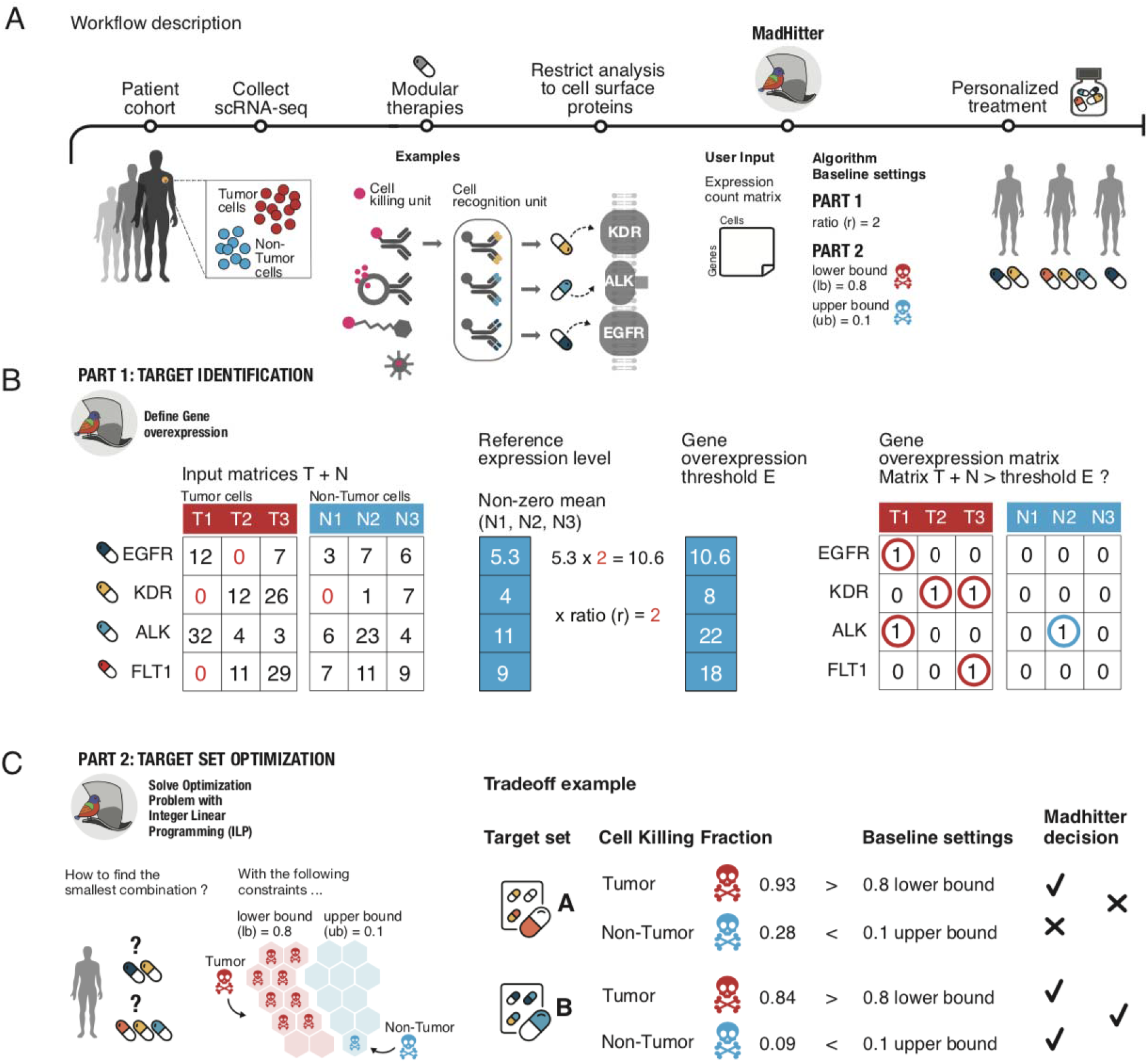
Conceptual schematic example of MadHitter analysis of single-cell data transcriptomics from three cancer patients. **(A)** A cohort of patients (three in this example) arrives for a study in which single-cell tumor microenvironment (TME) transcriptomics data are collected from each patient; the data are analyzed with MadHitter and each patient receives an optimal personalized combination of targeted therapies from a pre-specified set (pill bottle). MadHitter is aimed at optimizing combinations of targeted therapies that are *modular*, that is, having a recognition unit that is gene/protein-specific, and a joint killing subunit (similar for all gene targets). Icons of four such modular therapies are shown; we focus on the three for which the target protein must be on the cell surface and the two for which it must be a receptor, and we mention degraders only here. Three main algorithm parameters are denoted near the MadHitter icon in panel **A** and explained in the later panels. **(B)** The single-cell TME data are represented in two matrices with the genes as rows and cells as columns, partitioned into tumor (T) and non-tumor (N) cells. The expression ratio *r* determines by how much a gene must be overexpressed for a cell to be considered as a targeted. A gene is considered ‘overexpressed’ in either a non-tumor cell or a tumor cell if its expression is at least *r* times the mean, reference level; e.g, the reference level for *FLT1* is (7+11+9)/3 = 9 and only cell T3 has *FLT1* expression above 9×2 = 18. The matrices on the right side show a Boolean representation of which targets kill which cells, based on the expression values presented in this toy problem in matrix B and taking *r*=2. Accordingly, the combination of EGFR and KDR would kill all tumor cells and would spare all non-tumor cells. See another example in **Supplementary Figure S1 (C)** The main algorithm in MadHitter seeks a combination of targets that is as small as possible and would kill many tumor cells and few non-tumor cells, in a patient-specific manner. The *lb* and *ub* parameters are the lower bound on the fraction of tumor cells killed and the upper bound on the fraction of non-tumor cells whose killing is tolerated, respectively. Baseline settings used in our analyses are *r* = 2, *lb* = 0.8 and *ub* = 0.1, and are varied in some of the analyses. The right side of the panel shows a hypothetical example of the tradeoff between killing tumor cells and sparing non-tumor cells. While target set A could kill a larger fraction of tumor cells than target set B, MadHitter would select target set B since only it satisfies both our baseline settings and kills at most 0.1 fraction of the non-tumor cells.

## Results

### The Data and the Combinatorial Optimization Framework

We focused our analysis searching for optimal treatment combinations in nine single-cell RNAseq data sets that include tumor cells and non-tumor cells from at least three patients that were publicly available at the onset of our investigation (**Methods; Table 1**). Those data sets include four brain cancer data sets and one each from head and neck, melanoma, lung, breast and colon cancers. Most analyses were done for all data sets, but for clarity of exposition, we focused the main text analyses on four data sets from four different cancer types (brain, head and neck, melanoma, lung) that are larger than the other five and hence, make the optimization problems more challenging. The results for the other five data sets are provided in the **Supplementary Materials**.

**Table 1.** Summary descriptions of single-cell data sets from solid tumors used either for analysis (9) or preliminary testing (5 additional) and one liquid tumor data set used for validation (1 additional). Data sets are ordered so that those from the same or similar tumor types are on consecutive rows. The first 13 and 15th data sets were obtained either from GEO or the Broad Institute Single Cell Portal, but the GEO code is shown. The data set on the 14th row was obtained from ArrayExpress. In some data sets that have both cancer and non-cancer cells, there may be patients for whom only one type or the other is provided. Hence, the numbers in parentheses in the third and fourth columns may differ. In most cases, the number of patients we analyzed is the minimum of the two numbers in parentheses; for example, for GSE103322, we analyzed data on 13 patients, which is the minimum of (13) and (15). Data set GSE115978^64^ supersedes and partly subsumes GSE7256^65^. For the blood cancer data set, we used only the six samples from B-cell acute lymphoblastic leukemia (ALL) patients and report only those cells as counted in the version at (http://tisch.comp-genomics.org); the data set also contains samples from two patients with T-cell ALL and three healthy donors that we did not analyze.

| Data set code | Cancer type(s) | Cancer cells(patients) | Non-cancer cells (patients) | Clinical follow-up | Reference(s) |
| --- | --- | --- | --- | --- | --- |
| GSE75688 | Breast | 441(11) | -- | Metastasis or not | 66 |
| GSE118389 | Breast | 804(6) | 314(6) | Metastasis or not | 63 |
| GSE89567 | Brain (glioma) | 5097(10) | 1146(9) | No | 67 |
| GSE103224 | Brain (glioma) | 23793(8) | -- | No | 62 |
| GSE70630 | Brain (glioma) | 4044(6) | 303(6) | No | 68 |
| GSE57872 | Brain (glioma) | 440(6) | -- | No | 69 |
| GSE102130 | Brain (6 glioma and 3 glioblastoma) | 2858(9) | 94(5) | No | 70 |
| GSE84465 | Brain (glioblastoma) | 1091(4) | 651(4) | No | 45 |
| GSE81861 | Colorectal | 272(10) | 160(6) | No | 71 |
| GSE103322 | Head and Neck | 2093(13) | 3197(15) | No | 72 |
| GSE115978 | Melanoma | 2018(23) | 4334(32) | Yes, immuno-therapy | 64,65 |
| GSE118828 | Ovarian | 1415(11 primary)<br>973 (5 metastasis) | 578(2) | No | 73 |
| GSE67980 | Prostate | 124(21) | -- | Metastasis or not | 74 |
| E-MTAB-6149 | Lung | 7351(5) | 2730(5) | No | 75 |
| GSE132509 | Acute Lymphoblastic Leukemia (ALL) | 21329 (6 B-ALL) | 4744 (6 B-ALL) | No | 37 |

To formalize our questions as combinatorial optimization hitting set problems, we define the following parameters and baseline values and explore how the optimal answers vary as functions of these parameters: We specify a lower bound on the fraction of tumor cells that should be killed, *lb*, which ranges from 0 to 1. Similarly, we define an upper bound on the fraction of non-tumor cells killed, *ub*, which also ranges from 0 to 1. To be concrete, our baseline settings are *lb* = 0.8 and *ub* = 0.1 but we also explore other values, and the approach (and code) are generic and can explore the landscape of solutions at any other settings deemed of interest. To represent the concept that only cells that overexpress the target, we introduce an additional parameter *r*. The expression ratio *r* defines which cells are killed, as follows (**Figure 1B**): Denote the mean expression of a gene *g* in *non-cancer* cells that have non-zero expression by E(*g*). A given cell is considered killed if gene *g* is targeted and its expression level in that cell is at least *r* × *E*(*g*). Higher values of *r* thus model more selective killing.

Having *r* as a modifiable parameter anticipates that in the future one could experimentally tune the overexpression level (**see Supplementary Materials 1**) at which cell killing occurs^21^. In this respect, technologies that rely on RME to get a toxin into the cell are particularly tunable because there is known to be a non-linear relationship, called “*superselectivity*” between the number of protein copies on the cell surface and the probability that RME occurs successfully^34^. In these technologies, the toxin or other therapy delivered by the modular treatment enters cells in a gene-specific manner^35^.CAR-T therapy activates T-cell killing against cells in a gene-specific manner^21,22^.

The question of how sharp a non-linear superselective jump in ligand-receptor binding as a function of surface density of receptors is achievable in a nanoparticle system has been the focus of both experimental and theoretical studies for nanoparticles and CAR-T, as reviewed in **Supplementary Materials 1**. The inverse of the slope of this binding function at the transition is one way to estimate a realistic value of *r*. For nanoparticles, an increase of a factor of at most 2 in receptor density/expression appears to be sufficient to go from almost no binding to almost perfect binding and this justifies our baseline choice of *r* = 2.0. For CAR-T, investigation of the binding curve has only recently started, and the first key study achieved experimentally a receptor density ratio of 5-10 between almost no binding and almost perfect binding. These CAR-T experiments were a proof-of-principle study using combinatorial logic encoded genetically and it seems likely that more sophisticated CAR-T circuitry can achieve steeper binding curves corresponding to lower values of *r*. Therefore, for most of our analyses, the expression ratio *r* is varied from 1.5 to 3.0, with a baseline of 2.0, based on experiments in the lab of N.A. and related to combinatorial chemistry modeling^34^; in one analysis, we varied *r* up to 5.0 (**Supplementary Materials**).

Given these definitions, we solve the following combinatorial optimization hitting set problem (**Methods**): Given an input of a single-cell transcriptomics sample of non-tumor and tumor cells for each patient in a cohort of multiple patients, bounds *ub* and *lb*, ratio *r*, and a set of target genes, we seek a solution that includes a minimum-size combination of targets in each individual patient, while additionally minimizing the size of all targets given to the patient cohort. The latter is termed the *global minimum-size hitting set (GHS*) in computer science terminology or the *cohort target set (CTS)* in terminology specific to our problem, while the optimal hitting set of genes targeting one patient is termed the *individual target set* (*ITS*). This optimum hitting set problem with constraints *can be solved to optimality using integer linear programming (ILP)* (**Methods, Supplementary Figure S2**). We solve different optimization problem instances, each of which considers a different set of candidate target genes: 1269 genes encoding cell surface receptor proteins, and subset of 58 out of these 1269 genes that already have published ligand-mimicking peptides, and a nested collection of sets of 424-900 out of the 1269 genes that are lowly expressed across normal human tissues below a series of decreasing gene expression thresholds^22^. From a computational standpoint, there is no inherent theoretical limit on the size of the candidate gene set, but in practice, such optimization tasks may become intractable as this size increases. Our formulation is personalized as each patient receives the minimum possible number of treatments. The global optimization comes into play only when there are multiple solutions of the same size to treat a patient. For example, suppose we have two patients such that patient A could be treated by targeting either {*EGFR, FGFR2*} or {*MET, FGFR2*} and patient B could be treated by targeting either {*EGFR, CD44*} or {*ANPEP, CD44*}. Then we prefer the CTS {*EGFR, FGFR2, CD44*} of size 3 and we treat patient A by targeting {*EGFR, FGFR2*} and patient B by targeting {*EGFR, CD44*}.

As the number of cells per patient varies by three orders of magnitude across data sets, we use random sampling to obtain hitting set instances of comparable sizes that adequately capture tumor heterogeneity. We found that sampling hundreds of cells from the tumor is sufficient to get enough data to represent all cells. In most of the experiments shown, the number of cells sampled, which we denote by *c*, was 500. In some smaller data sets, we had to sample smaller numbers of cells (**Methods**). As shown in (**Supplementary Materials 2**, **Figures S2-S3**), 500 cells, when available, are roughly sufficient for CTS size to plateau for our baseline parameter settings, *lb* = 0.8, *ub* = 0.1, *r* = 2.0. The results using all cells and default parameters are shown in **Supplementary Table S1** and are similar to the results using sampling, where we consider the latter more informative for future studies. Hence, for each individual within a data set, we performed independent sampling of *c* cells 20 times and their results were summarized.

### Cohort and Individual Target Set Sizes as Functions of Tumor Killing and Non-Tumor Sparing Goals

Given the single-cell tumor data sets and the ILP optimization framework described above, we first studied how the resulting optimal cohort target set (CTS) may vary as a function of the parameters defining the optimization objectives in different cancer types. **Figures 2 and S4-S8 in Supplementary Materials 3** show heatmaps of CTS sizes when varying *lb*, *ub*, and *r* around the baseline values of 0.8, 0.1, and 2.0, respectively. The CTS sizes for melanoma were largest, partly due to the larger number of patients in that data set (**Table 1**). Indeed, as we sampled subsets of 5 or 10 patients uniformly and observed that the mean CTS sizes grew from 7.9 (5 patient subsets) to 12.3 (10 patient subsets) to 31.0 (all patients, as shown in **Figure 2**).

**Figure 2.**
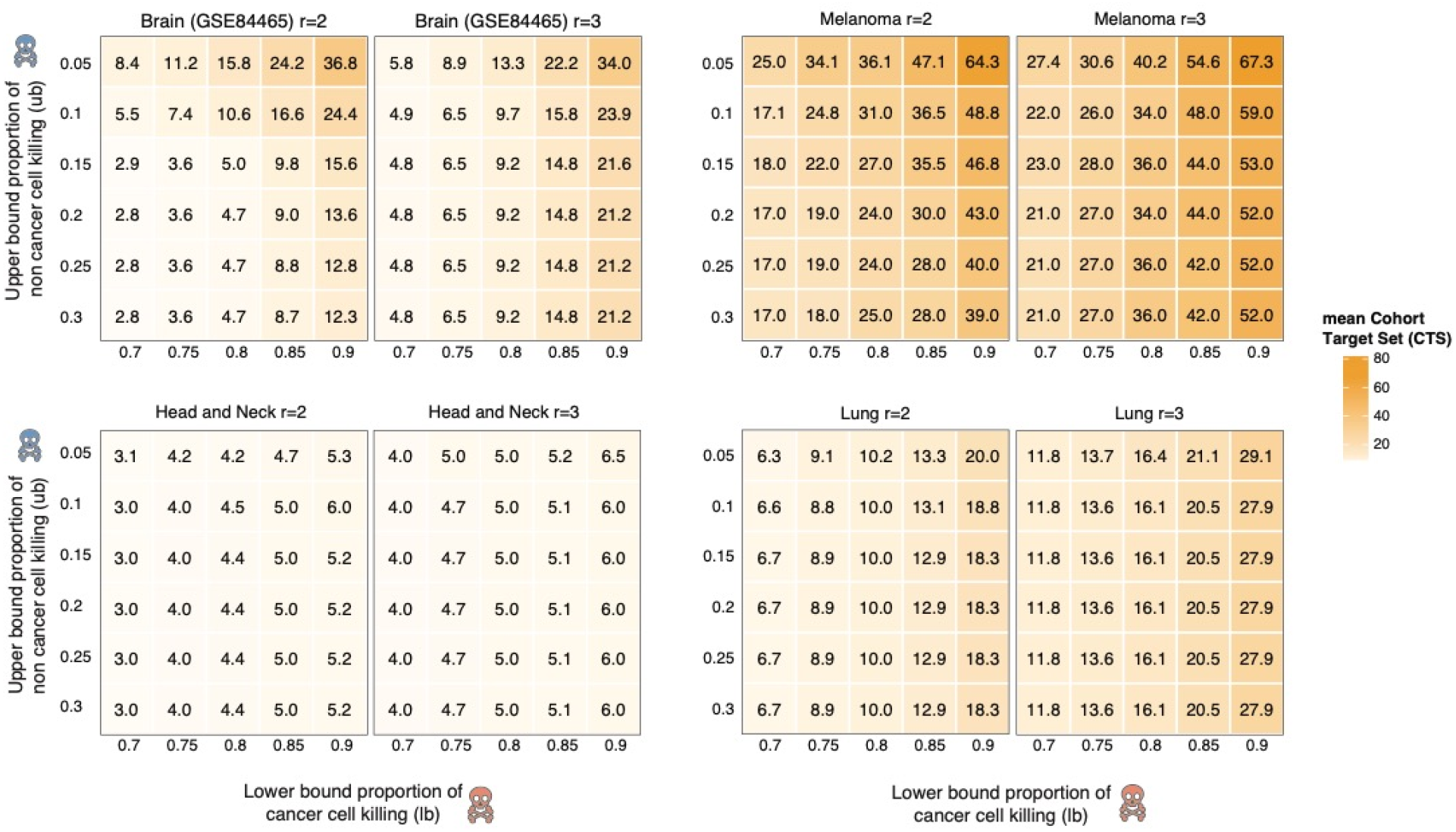
Heat maps showing how the cohort target set size (CTS) varies as a function of *lb, ub, r* and across data sets. For each plot the x-axis and y-axis represent *lb* and *ub* parameter values, respectively. The scale on the right shows the cohort target set sizes by color scale. We show separate plots for *r* = 2.0, 3.0 here and a larger set {1.5, 2.0, 2.5, 3.0} in **Supplementary Materials 3**. Individual values are not necessarily integers because each value represents the mean of 20 replicates of sampling *c* (500 for each of the data sets shown here) cells (**Figure S2**).

Encouragingly, for most data sets and parameter settings, the optimal CTS sizes are in the single digits. However, in several data sets, we observe a sharp increase in CTS size as *lb* values are increased above 0.8 and/or as the *ub* is decreased below 0.1, with a more pronounced effect of varying *lb*. This transition is more discernable at the lowest value of *r* (1.5), probably because when *r* is lower, it becomes harder to find genes that are individually selective in killing tumor cells and sparing non-tumor cells (**Supplementary Figures S4-S8**). The qualitative transition observed in CTS sizes occurs robustly regardless of the threshold for filtering out low expressing cells when preprocessing the data (**Supplementary Materials 4, Figures S9-S11**).

We next examined what are the resulting *individual target set* (ITS) sizes obtained in the optimal combinations under the same conditions. In all data sets, the mean ITS sizes are in the single digits for most values of *lb* and *ub*. The distributions of ITS sizes are shown for four data sets and two combinations of (*lb, ub*) (**Figure 3**) and for additional data sets in **Supplementary Materials 5, Figure S12**. Overall, the mean ITS sizes with the baseline parameter values (*r* = 2.0, *lb* = 0.8, *ub* - 0.1) range from 1.0 to 3.91 among the nine data sets studied (**Supplementary Table S3**); on average 4 targets per patient should hence suffice if enough single-target treatments are available in the cohort target set. However, there is considerable variability across patients.

**Figure 3.**
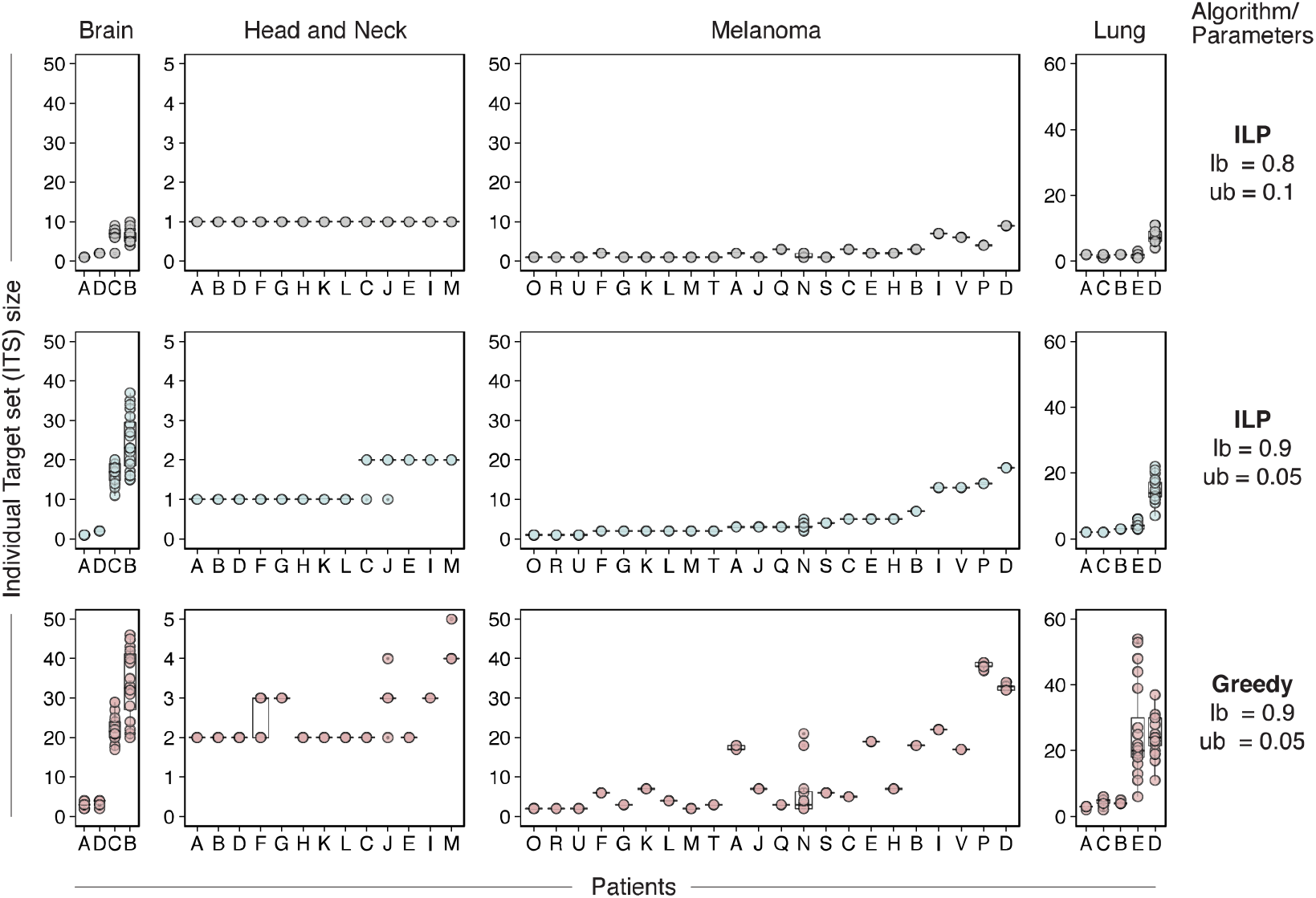
The distribution of optimal and greedy individual treatment combination sizes (ITS) values in four different cancer types. We study both our baseline parameter setting (upper row panels) and a markedly more stringent one (middle row plots). For the more stringent parameter setting, we compare the ITS sizes obtained using MadHitter (middle row plots) and a greedy algorithm that tries to add pairs of genes at a time (bottom row plots). In each plot, the patients are sorted from left to right according to their mean ITS values in the optimal stringent regime. Additional comparisons between ITS sizes at different parameter settings can be found in **Supplementary Materials 5**. Description of the greedy algorithm and more comparisons between the optimal and greedy algorithms are provided in **Supplementary Materials 8**.

Evidently, as we make the treatment requirements more stringent (by increasing *lb* from 0.8 to 0.9 and decreasing *ub* from 0.1 to 0.05), the variability in ITS size across patients became larger. Importantly, this analysis provides rigorous quantifiable evidence grounding the prevailing observation that among tumors of the same type, some individual tumors may be much harder to treat than others. Taken together, these results show that we can compute precise estimates of the number of targets needed for cohorts (in the tens) and individual patients (in the single digits usually) and that these estimates are sensitive to the killing stringency, especially when the *lb* increases above 0.8. The variation for more aggressive killing regimes, with values of *lb* up to 0.99 for the baseline *r* = 2.0 is displayed in **Figures S13-S14 in Supplementary Materials 6**. For fixed *lb* = 0.8, *ub* = 0.1 and varying *r*, to values as high as 5.0, smallest CTS sizes are typically obtained for *r* values close to 2.0, further motivating our choice of *r* = 2 as the default value (**Supplementary Materials 7, Figures S15-S16, Supplementary Table S2**). Finally, we show that, as expected, a ‘control’ greedy heuristic algorithm searching for small and effective target combinations finds ITS sizes substantially larger than the optimal ITS sizes identified using our optimization algorithm (**Figure 3**). The greedy CTS size is greater than the ILP optimal CTS size for eight out of nine data sets (**Table S3** in **Supplementary Materials 8, Methods**). The comparison between heuristic and optimal solutions is of interest because it quantifies the benefit of finding the optimum-size ITS and CTS and the previous related studies reviewed in the Introduction used only heuristic methods for solution sizes above 2 or 3 respectively^22,35^.

### The Landscape of Combinations Achievable with Receptors Currently Targetable by Published Ligand-Mimicking Peptides or those Tested in CAR-T Trials

To get a view of the combination treatments that are possible with receptor targets for which there are already existing modular targeting reagents, we conducted a literature search identifying 58 out of the 1269 genes with published ligand-mimicking peptides that have been already tested in *in vitro* models, usually cancer models (**Methods**; **Tables 3 and 4**). We asked whether we could find feasible optimal combinations in this case and if so, how do the optimal CTS and ITS sizes compare vs. those computed for all 1269 genes?

Computing the optimal CTS and ITS solutions for this basket of 58 targets, we found feasible solutions for six of the data sets across all parameter combinations we surveyed and three of these six are illustrated for each patient in **Figure 4**. However, for three data sets, in numerous parameter combinations we could not find optimal solutions that satisfy the optimization constraints (**Supplementary Materials 9, Figures S17-S19**). That is, the currently available targets do not allow one to design treatments that may achieve the specified selective killing objectives, underscoring the need to develop new targeted cancer therapies, to make personalized medicine more effective for more patients.

**Figure 4.**
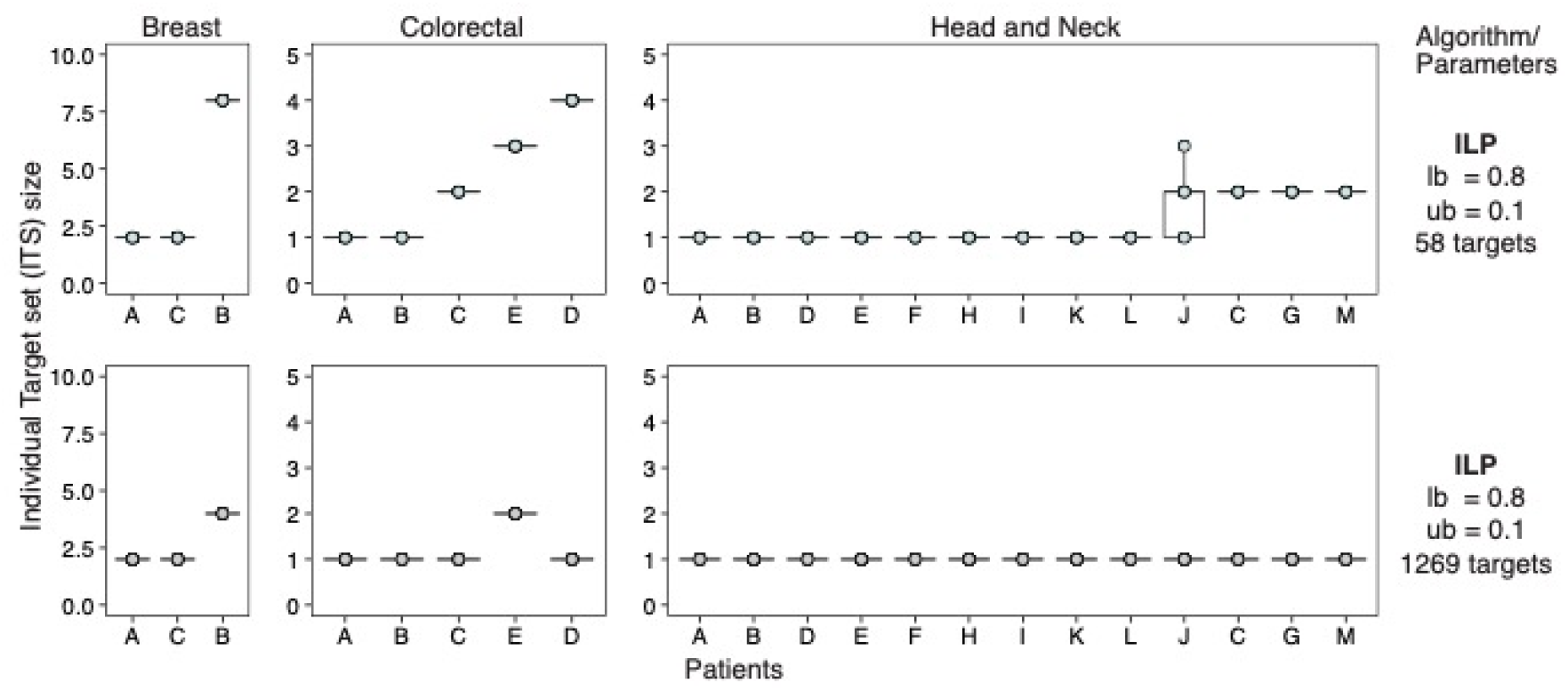
Comparison of Individual Target Set Sizes with 1269 or 58 targets for three out of the six data sets that have feasible solutions. We attempted to find feasible solutions for all patients using 58 cell surface receptors that have published ligand-mimicking peptides that have been tested *in vitro* or in pre-clinical models. There are feasible solutions for all patients in six data sets, but not for the brain (GSE84465), melanoma (GSE115978), and lung (E-MTAB-6149), which were displayed in previous figures. Instead, we show here results for breast and colorectal cancers, for which other analyses, such as those in **Figures 2** and **3**, are in the **Supplementary Materials.** Some of the optimal solutions obtained on the 58-receptors restricted set are of the same size to those obtained on the whole receptors set and some are larger.

Overall, comparing the optimal solutions obtained with 58 targets to those we have obtained with the 1269 targets, three qualitatively different behaviors are observed (**Supplementary Materials 9, Figures S17-S19**): (1) In some datasets, it is just a little bit more difficult to find optimal ITS and CTS solutions with the 58-gene pool, while in others, the restriction to a smaller pool can be a severe constraint making the optimization problem infeasible. (2) The smaller basket of gene targets may force more patients to receive similar individual treatment sets and thereby reduces the size of the CTS. (3) Unlike the CTS size, the ITS size must stay the same or increase when the pool of genes is reduced, because we find the optimal ITS size for each patient. Overall, the average ITS sizes across each cohort using the pool of 58 genes for baseline settings range from 1.16 to 4.0. Among cases that have any solution, the average increases in the ITS sizes at baseline settings in the 58 genes case vs. that of the 1269 case were moderate, ranging from 0.16 to 1.33.

We performed a similar analysis with 57 gene targets that were listed as the genes encoding single proteins in CAR-T trials (**Supplementary Table S4;** see one visualization of the results in **Supplementary Figure S20)**^22^. However, using our baseline parameter settings and either with cell sampling or analyzing all cells together only three out of nine data sets have feasible solutions because the majority of the CAR-T target genes were filtered out by the researchers who collected the primary data. Those include brain cancer GSE89567 (mean CTS 2.95, ITS 1), brain GSE102130 (mean CTS 4, ITS 2) and breast cancer GSE118389 (mean CTS 7, ITS 3). This limited success suggests that, to cover more cancer indications, CAR-T targets that are more differentially expressed in solid tumors need to be developed and that it may be necessary to target multiple receptors simultaneously^22,23^.

The success of CAR-T targeted to CD19 for B-cell acute lymphocytic leukemia (B-ALL)^36^ and the availability of one single-cell B-ALL data set (GSE132509, **Methods**)^37^ allowed us to assess further whether our 0.8 killing threshold for malignant cells reasonably captures the success observed in a human trial. After filtering out cells expressing less than 20% of genes and using only the cells consistently annotated as malignant in the six B-ALL samples, we found that a mean of 80.3% per patient of malignant cells express CD19 (**Methods**). This result is consistent with the 0.8 baseline value of *lb* we have used throughout the paper. The annotations of the non-malignant cells were not sufficiently consistent to assess the 0.1 baseline value of *ub*. Notably, blood cancers are fundamentally different from solid tumors. A different line of support for the choice of the 0.8 threshold is provided in **Methods**.

### Optimal Fairness-Based Combination Therapies for a Given Cohort of Patients

Until now we have adhered to a *patient-centered approach* that aims to find the minimum-size ITS for each patient, first and foremost. We now study a different, *cohort-centered approach*, where given a cohort of patients, we seek to minimize the total size of the overall CTS size, while allowing for some increase in the ITS sizes. The key question is how much larger are the resulting ITS sizes if we optimize for minimizing the cohort (CTS size), rather than the individuals (ITS size)? This challenge is motivated by a ‘fairness’ perspective (**Supplementary Materials 1**), where we seek solutions that are beneficial for the entire community from a social or economic perspective (in terms of CTS size) even if they are potentially sub-optimal at the individual level (in terms of ITS sizes). Here, the potential benefit is economic since running a basket trial would be less expensive if one reduces the size of the basket of available treatments (**Figure 5A-B**).

**Figure 5.**
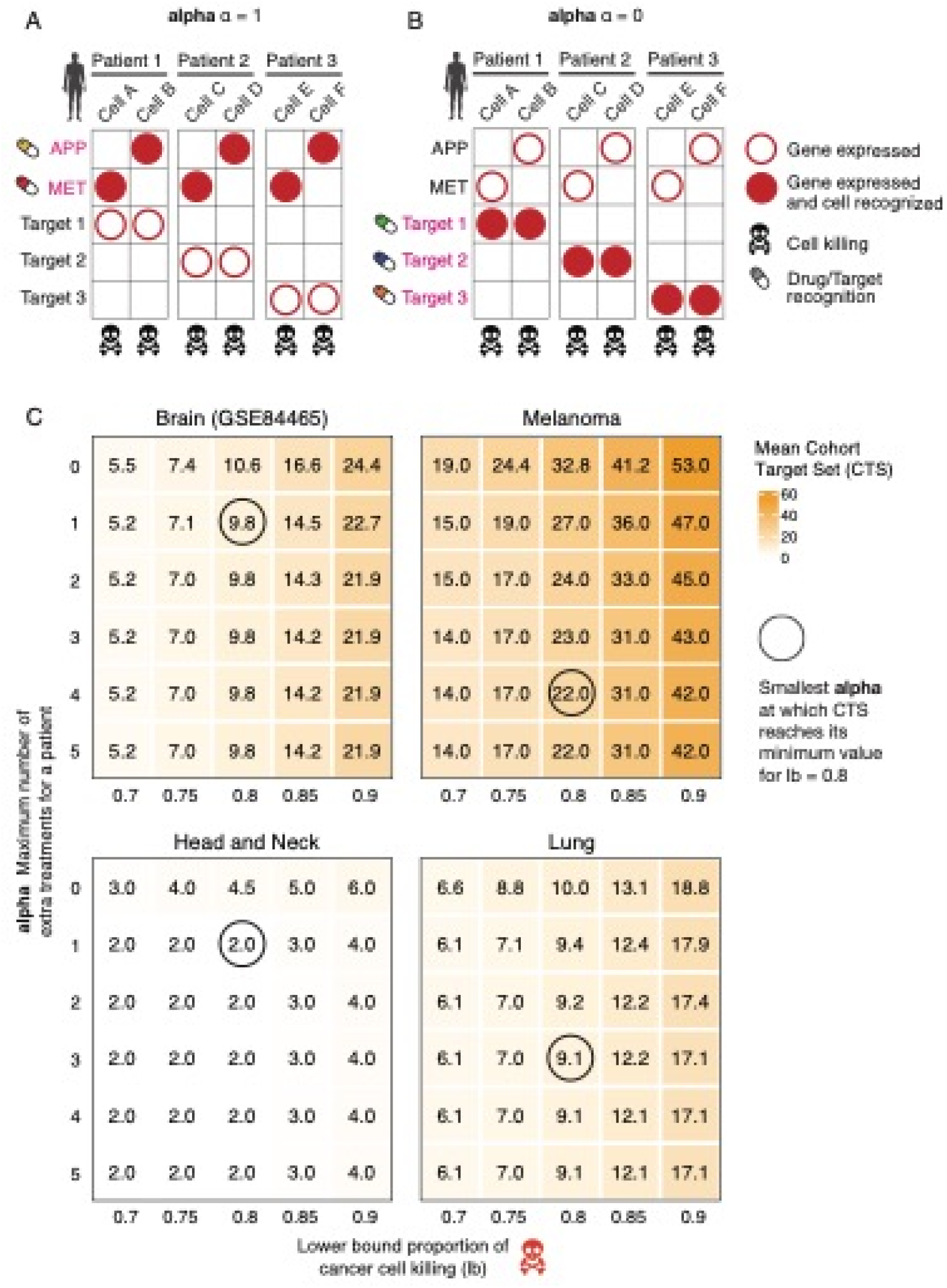
A schematic example demonstrating the rationale and workings of *fairness*-based solutions. **(A, B)** Let us assume that each of three patients has two tumor cells (columns), each displaying five membrane receptors that are highly expressed only on the tumor cells and not on the non-tumor ones (rows). If we target {APP, MET} (panel **A,** *α* = 1) in all patients, then this achieves a CTS size of 2, which is the minimum possible. Employing the original individual-based optimizing objective, each patient could instead be treated by an ITS of size 1 by targeting the distinct receptors called Target 1 (specific to Patient 1), Target 2 and Target 3, respectively, but this would result in an optimal CTS of size 3 (panel **B,** *α* = 0). The solution in panel A has an unfairness value *α* = 1 because the worst difference among all patients is that a patient receives 1 more treatment than necessary. (**C**) Heatmaps showing how the CTS size varies as *α* increases (y-axis), starting from its baseline value of 0 where each patient is assigned a minimum-sizes individual treatment set (top row). The lower bound on tumor cells killed (x-axis) is also varied while the upper bound on non-tumor cells killed is kept fixed at 0.1. We are particularly interested in finding the smallest value on the y-axis at which the CTS size reaches its minimum value, which is circled for the baseline *lb* = 0.8, because this bounds the tradeoff between the achievable reduction in the number of targets needed to treat the whole cohort and the number of extra targets above the ITS minimum that any patient might need to receive.

We formalized this ‘fair CTS problem’ by adding a *cost parameter α* that specifies the limit on the excess number of (ITS) targets selected for any individual patient, compared to the number selected in the individual-based approach that was studied up until now (formally, the latter corresponds to setting *α* = 0). We formulated and solved via ILP this fair CTS problem for up to 1269 possible targets on all nine data sets (**Methods**). We fixed *r* = 2 and *ub* = 0.1 while varying *α* and *lb*. **Figure 5C** and **Figures S21-S25** in **Supplementary Materials 10** show the optimal CTS and ITS sizes for *α* = 0,...,5.

For 8 out of 9 data sets, we encouragingly find that the unfairness cost parameter *α is bounded by a constant of 3;* i.e., it is sufficient to increase *α* by no more than 3 to obtain the smallest CTS sizes in the optimally fair solutions. For the largest data set (melanoma), *α* = 4. As we show in **Supplementary Materials 10**, empirically, even if one requires lower α values, then as those approach 0, the size of the fairness-based CTS grows fairly moderately and remains in the lower double digits, and the mean size of the number of treatments given to each patient (their ITS) is overall < 5. Theoretically, we show that one can design instances for which *α* would need to be at least 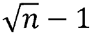 to get a CTS of size less than the overall number of targets *n* (**Supplementary Materials 10**). However, in practice, we find that given the current tumor single-cell expression data, fairness-based treatment strategies are likely to be a reasonable economic option in the future.

### The Landscape of Optimal Solutions Targeting Differentially Expressed Tumor Receptors that are Lowly Expressed Across Many Healthy Tissues

We turn to examine the space of optimal solutions when restricting the set of eligible surface receptor gene targets to those that have lower expression across many noncancerous human tissues (**Methods**), aiming to mitigate potential damage to tissues unrelated to the tumor site. To this end, we selected subsets of the 1269 cell surface receptor targets in which the genes have overall lower expression across multiple normal tissues, by mining GTEx and the Human Protein Atlas (HPA) (**Methods**). Varying the *selectivity expression thresholds* (expressed in transcripts per million (TPM)) used to filter out genes whose mean expression across the normal adult tissues is above values of 10, 5, 2, 1, 0.5, and 0.25 (i.e., employing more and more extensive filtering as this threshold is decreased), decreases the size of the target cell surface receptor gene list by more than half (**Table 2**).

**Table 2:** Size of high confidence target gene sets for different thresholds.

| Thresholds<br>expression across | Size of gene set | No of genes which<br>are NOT olfactory | No of genes with<br>ligand mimicking |
| --- | --- | --- | --- |

| tissues (TPM) |  | (OR*) and taste receptors (TAS*) | peptides (intersection with Tables 3 and 4) |
| --- | --- | --- | --- |
| 0.25 | 424 | 97 | 1 |
| 0.5 | 494 | 141 | 3 |
| 1 | 547 | 187 | 3 |
| 2 | 632 | 269 | 6 |
| 5 | 762 | 398 | 10 |
| 10 | 900 | 536 | 19 |

As shown in **Figures 6A, B (and Supplementary Figures S26-S28)**, MadHitter identifies very different cohort target sets (which are larger than the original optimal solutions, as expected) as the TPM selectivity threshold value is decreased. Furthermore, different ITS instances may become infeasible (**Supplementary Figure S29**). At an individual patient level, using lower selectivity threshold levels, which leads to a smaller space of membrane receptors to choose from, also leads to increased mean ITS sizes (**Supplementary Figures S30, S31**). Across the nine data sets, the selectivity threshold at which the CTS problem became infeasible varied (**Supplementary Figure S29**). The differences observed could be the result of expression heterogeneity of the cancer, number of patients within the data set, size of target gene set, lack of expression of available gene targets and other unknown factors. In the future, further experimentation is required to identify tissue-specific optimal gene expression thresholds that will minimize side effects while allowing cancer cells to be killed by combinations of targeted therapies. Finally, for completeness, we also tested MadHitter on the set of 533 lowly expressed genes suggested by MacKay et al.^25^ All instances with default setting of *r, lb, ub* have feasible solutions for all patients. Mean ITS sizes are below 4 for eight of nine data sets, but close to 10 for the brain cancer data set GSE84465. More details can be found in **Supplementary Materials 11** and **Table S5**.

**Figure 6.**
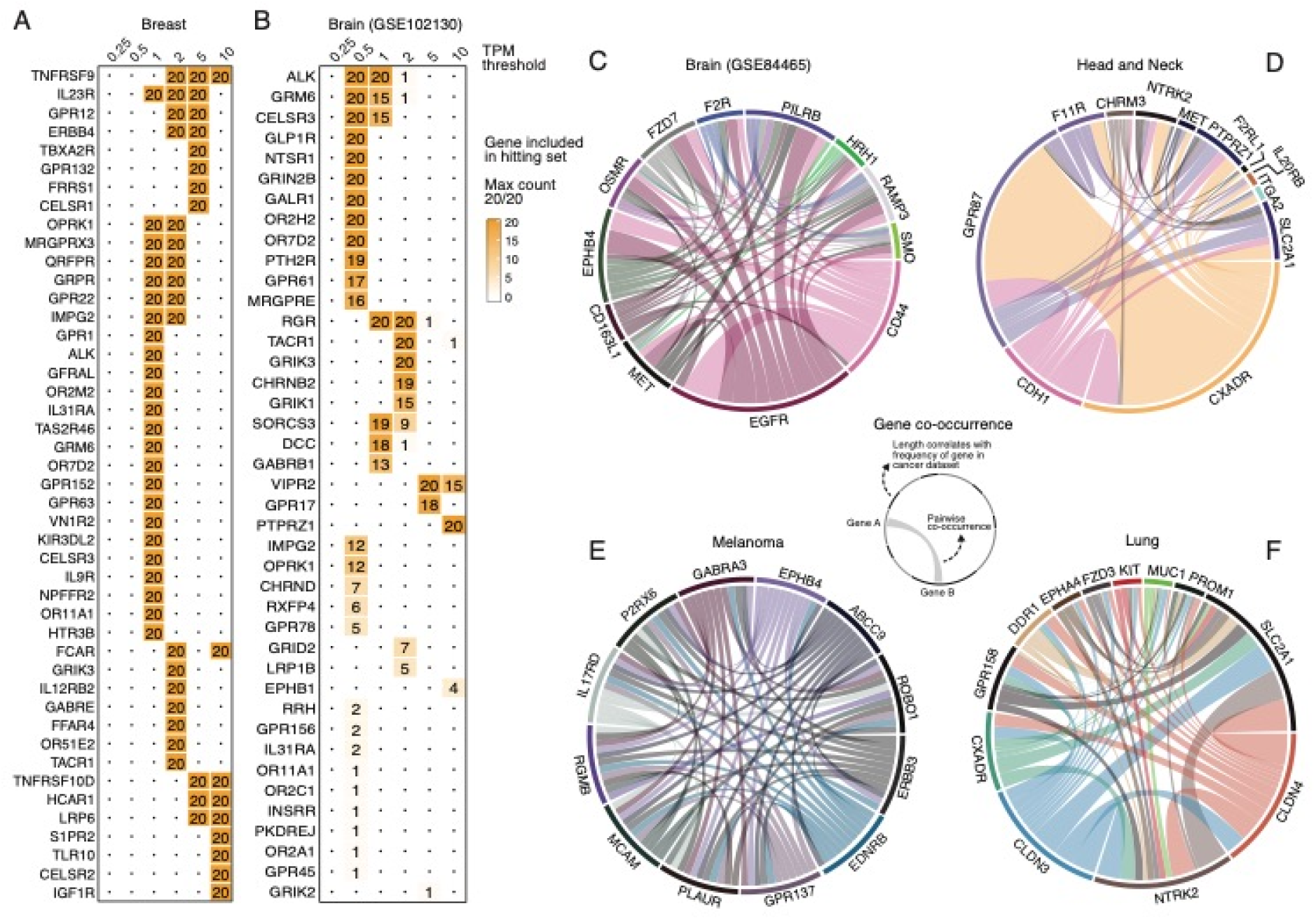
Variation in the CTS size and composition as function of the magnitude of filtering of genes expressed in noncancerous human tissues, for different tumor types. **(A-B)** The number of times a gene (cell-surface receptor) is included in the CTS (out of 20 replicates, which is therefore the Max count in panels A-B), where each column presents the CTS solutions when the input target genes sets are filtered using a specific TPM filtering threshold (**Methods**), for (**A)** a breast cancer and **(B)** brain cancer. These data sets were selected due to their relatively small cohort target set sizes, permitting their visualization. **(C-F)** Circos plots of the genes occurring most frequently in optimal CTS solutions (length of arc along the circumference) and their pairwise co-occurrence (thickness of the connecting edge) for the four main cancer types, in our original target space of 1269 encoding cell-surface receptors. For each data set, we sampled up to 50 optimal CTS solutions. Co-occurrence representations of the 12 most common target genes out of 1269 encoding cell-surface receptors (with greater than 5% frequency of occurrence) are represented in a cancer specific manner for **(C)** brain cancer, **(D)** head and neck cancer, (only seven genes have a frequency of 5% or more across optimal solutions), **(E)** melanoma, and **(F)** lung cancer. Genes and connections have distinct colors for improved visibility.

### Key Targets Composing Optimal Solutions Across the Space of 1269 membrane receptors

To identify the genes that occur most often in optimal solutions for our baseline settings, since there may be multiple distinct optimal solutions composed of different target genes, we sampled up to 50 optimal solutions for each optimization instance solved and recorded how often each gene occurs and how often each pair of genes occur together (**Methods**). We analyzed and visualized these gene (co-)occurrences in three ways. First, we constructed co-occurrence circos plots in which arcs around the circle represent frequently occurring genes and edges connect targets that frequently co-occur in optimal CTS solutions. **Figure 6C-F** shows the co-occurrence visualizations for optimal CTS solutions obtained with the original, unfiltered target space of 1269 genes and in baseline parameter settings. The genes frequently occurring in optimal solutions are quite specific and distinct between different cancer types. In melanoma, the edges form a clique-like network because virtually all optimal solutions include the same clique of 12 genes (**Figure 6E**). The head and neck cancer data set has only one commonly co-occurring pair {*GPR87, CXADR*}, partly because the CTS sizes are much smaller, ranging from 2 to 5, and the optimal solutions of sizes larger than 2 vary substantially in the choices of the other gene(s) (**Figure 6D**). The choices of these two genes are not obvious as *CXADR* ranks only 10^th^ by our measure of gene overexpression (**Supplementary Materials 8**) and *GPR87* is not in the top 20 genes. Of the cancer types not depicted in **Figure 6**, the breast cancer data set has a commonly co-occurring set of size 4, {*CLDN4, INSR, P2RY8, SORL*} none of which ranks in the top 20 genes, and the colorectal cancer data set has a different commonly co-occurring set of size 4, {*GABRE, GPRR, LGR5, PTPRJ*} (data not shown). These variations should be viewed as characteristic of different data sets, not of different tumor types, since we see different results for the four brain cancer data sets.

We next tabulated sums of how often each gene occurred in optimal solutions for all nine data sets (**Supplementary Materials 12, Tables S6, S7 and S8**), obtained when solving for either 58 gene targets or 1269 gene targets. Strikingly, one gene, *PTPRZ1* (protein tyrosine phosphatase receptor zeta 1), appears far more frequently than others, especially in three brain cancer data sets (GSE70630, GSE89567, GSE102130, **Supplementary Table S8**). *PTPRZ1* also occurs commonly in optimal solutions for the head and neck cancer data set (**Figure 6D**). The brain cancer finding coincides with previous reports that *PTPRZ1* is overexpressed in glioblastoma (GBM)^38,39^. *PTPRZ1* also forms a fusion with the nearby oncogene *MET* in some brain tumors that have an overexpression of the fused *MET*^40^. Notably, various cell line studies and mouse studies have shown that inhibiting PTPRZ1, for example by shRNAs, can slow glioblastoma tumor growth and migration^41,42^. There have been some attempts to inhibit PTPRZ1 pharmacologically in brain cancer and other brain disorders^43,44^. In the four brain cancer data sets, *PTPRZ1* is expressed selectively above the baseline *r* = 2.0 in 0.99 (GSE89567), 0.84 (GSE70630), 0.96 (GSE102130) and 0.27 (GSE84465) proportion of cells in each cohort. The much lower relative level of *PTPRZ1* expression in GSE84465 is likely due to the heterogeneity of brain cancer types in this data set^45^. Among the 58 genes with known ligand-mimicking peptides, *EGFR* stands out as most common in optimal solutions (**Supplementary Table S6**). Even when all 1269 genes are available, *EGFR* is most commonly selected for the brain cancer data set (GSE84465) in which *PTPRZ1* is not as highly overexpressed (**Figure 6C**).

*PTPRZ1* was the fifth most frequently occurring gene in optimal solutions for the head and neck cancer data set (GSE103322). The two most common genes by a large margin are *CXADR* and *GPR87*. *CXADR* has been studied primarily by virologists and immunologists because it encodes a receptor for cocksackieviruses and adenoviruses^46^. In one breast cancer study, CXADR was found to play a role in regulating PTEN in the AKT pathway, but CXADR was underexpressed in breast cancer^47^ whereas it is overexpressed in the head and neck cancer data we analyzed. *GPR87* is a rarely studied G protein-coupled receptor with an unknown natural ligand^48^. In the context of cancer, GPR87 has previously been reported as overexpressed in several tumor types including lung and liver^48^ and its overexpression may play an oncogenic role via either the p53 pathway^49^ the NFκB pathway^50^ or other pathways.

Finally, we analyzed the set of genes in optimal solutions via the STRING database and associated tools^51^ to perform several types of gene set and pathway enrichment analyses. **Figures S32-S35 (Supplementary Materials 12)** show STRING-derived protein-protein interaction networks for the 25 most common genes in the same four data for which we showed co-occurrence graphs in **Figure 6C-F**. Again, *EGFR* stands out as being a highly connected protein node in the solution networks for both the brain cancer and head and neck cancer data sets. Among the 30 genes in the 1269-gene set that occur most commonly in optimal solutions (**Supplementary Table S7**), there are six kinases (out of 88 total human transmembrane kinases with a catalytic domain, STRING gene set enrichment *p* < 1*e* - 6), namely {*EGFR, EPHB4, ERBB3, FGFR1, INSR, NTRK2*} and two phosphatases {*PTPRJ, PTPRZ1*}. The KEGG pathways most significantly enriched, all at *FDR* < 0.005, are (“proteoglycans in cancer”) represented by {*CD44, EGFR, ERBB3, FGFR1, PLAUR*}, (“adherens junction”) represented by {*EGFR, FGFR1, INSR, PTPRJ*}, and (“calcium signaling pathway”) represented by {*EDNRB, EGFR, ERBB3, GRPR, P2RX6*}. The one gene in the intersection of all these pathways and functions is *EGFR*.

## Discussion

In this multi-disciplinary study, we harnessed techniques from combinatorial optimization to analyze publicly available single-cell tumor transcriptomics data to chart the landscape of future personalized combinations that are based on ‘modular’ therapies, including CAR-T therapy. We showed that, for most tumors we studied, four modular medications targeting different overexpressed receptors may suffice to selectively kill most tumor cells, while sparing most of the non-cancerous cells (**Figures 2** and **3** and **Table S3**. For the more restricted sets of low-expression genes^22^ or the 58 receptors with validated ligand-mimicking peptides (**Tables 3** and **4**), some patients do not have feasible solutions, especially as we reduce the TPM expression used for filtering the gene set to avoid targeting non-cancerous tissues. These findings indicate, on one hand, that researchers designing ligand-mimicking peptides have been astute in choosing targets relevant to cancer. On the other hand, these results suggest that there is a need for extending the set of cell surface receptors that can be targeted to enter tumor cells with ligated chemotherapy agents. There are two established methods to identify ligand-mimicking peptides called “one bead one compound” and “phage display”^20^, the latter of which was awarded the 2018 Nobel Prize in Chemistry. Both methods are applicable to all cell surface proteins and thus, we believe that the set of proteins with validated peptides could be greatly expanded as nanoparticle-based treatments get closer to being used in the clinic.

**Table 3.** Single proteins that can be targeted by peptides based on references 18, 19 and are expressed on the cell surface^78^. For easier correspondence with the gene expression data, the entries are listed in alphabetical order by gene symbol. In this table, we follow the clinical genetics formatting convention that proteins are in Roman and gene symbols are in *italics*.

| Protein | Gene Symbol |
| --- | --- |
| APN/CD13 | *ANPEP* |
| APP | *APP* |
| PD-L1 | *CD274* |
| CD44 | *CD44* |
| P32/gC1qR | *CD93* |
| E-cadherin | *CDH1* |
| N-cadherin | *CDH2* |
| CD21 | *CR2* |
| EGFR | *EGFR* |
| Epha2 | *EPHA2* |
| EphB4 | *EPHB4* |
| HER2 | *ERBB2* |
| FGFR1 | *FGFR1* |
| FGFR2 | *FGFR2* |
| FGFR3 | *FGFR3* |
| FGFR4 | *FGFR4* |
| VEGFR1 | *FLT1* |
| VEGFR3 | *FLT4* |
| PSMA | *FOLH1* |
| GPC3 | *GPC3* |
| IL-10RA | *IL10RA* |
| IL-11Rα | *IL11RA* |
| IL-13Rα2 | *IL13RA2* |
| IL-6Rα | *IL6R* |
| GP130 | *IL6ST* |
| VEGFR2 | *KDR* |
| MUC18 | *MCAM* |
| Met | *MET* |
| MMP9 | *MMP9* |
| Thomsen-Friedenreich carbohydrate antigen | *MUC1* |
| NRP-1 | *NRP1* |
| PDGFRβ | *PDGFRB* |
| CD133 | *PROM1* |
| PTPRJ | *PTPRJ* |
| HSPG | *SDC2* |
| E-selectin | *SELE* |
| Tie2 | *TEK* |
| VPAC1 | *VIPR1* |

**Table 4.** Single proteins that can be targeted by ligand-mimicking peptides but are not included in the two principal reviews that we consulted^19–20^ and are among 1269 cell surface receptors^78^. Since the evidence that these 20 genes have ligand-mimicking peptides is scattered in the literature, we include at least one PubMed ID of a paper describing a suitable peptide.

| Protein | Gene Symbol | At Least One PubMed ID |
| --- | --- | --- |
| ActRIIB | *ACVR2B* | 28955765 |
| CD163 | *CD163* | 27563889 |
| CXCR4 | *CXCR4* | 19482312, 22523575 |
| ephrin A4 | *EPHA4* | 15681844, 22523575 |
| ephrin B1 | *EPHB1* | 15722342, 22523575 |
| ephrin B2 | *EPHB2* | 15722342, 22523575 |
| ephrin B3 | *EPHB3* | 15722342, 22523575 |
| gonadotrophin releasing hormone receptor | *GNRHR* | 20814857, 22523575 |
| G Protein coupled receptor 55 | *GPR55* | 28029647 |
| bombesin receptor 2 | *GRPR* | 20814857, 22523575 |
| IL4 receptor | *IL4R* | 19012727 |
| low density lipoprotein receptor | *LDLR* | 27656777 |
| leptin receptor | *LEPR* | 19233229, 26265355 |
| LRP1 | *LRP1* | 29090274 |
| melanocortin 1 receptor | *MC1R* | 22964391 |
| melanocortin 4 receptor | *MC4R* | 17591746 |
| CD206 | *MRC1* | 30768279 |
| urokinase plasminogen activator receptor | *PLAUR* | 25080049 |
| neurokinin-1 receptor | *TACR1* | 29498264 |
| VPAC2 | *VIPR2* | 30077368 |

Remarkably, we found that if one designs the optimal set of treatments for an entire cohort adopting a fairness-based policy, then the size of the projected treatment combinations for individual patients are at most 3 targets larger, and in most data sets at most 1 target receptor larger than the optimal solutions that would have been assigned for these patients based on an individual-centric policy (**Figure 5, Supplementary Materials 10**). This suggests that the concern that the personalized treatment for any individual will be suboptimal solely because that individual happens to have registered for a cohort trial appears to be tightly bounded.

Like the study of MacKay et al.^22^, our study is a conceptual computational investigation. As in the studies of Dannenfelser et al.^21^ and MacKay et al.^22^, we assumed that treatment selection will be based on measuring gene expression; neither we nor they considered the possibility of treatments that target mutant proteins. Our framework could accommodate mutant proteins, if mutations are called from the single-cell mRNA reads, but this is challenging since the reads usually do not cover full mRNAs. We studied nine data single-cell expression data sets for the first time, but it would be helpful to analyze more and larger data sets in the future. Even among four data sets of the same (brain) cancer type, we observed considerable variability in CTS and ITS sizes. Since our approach is general and the software is freely available as source code, other researchers can test the method on new data sets or add new variables, constraints, and optimality criteria. For example, we chose to analyze each single cell as a unit; many analyses of scRNA cancer data choose to cluster the cells first and may instead use cell clusters as the units of analysis. In that view, analysis of differential gene expression is usually done between clusters, while we preferred to compare the expression of each gene in each cell to the expression of the same gene in all non-cancer cells. Our investigation may lead others to further improve our method and broaden its applicability. In future work, we plan to apply our approach to study ways for selectively killing specific populations of immune cells, such as myeloid-derived suppressor cells or T regulatory cells, because they inhibit tumor killing, while sparing most other non-cancer cells.

We compared gene expression levels between non-cancer cells and cancer cells sampled from the same patient, which avoids inter-patient expression variability^52^. Following recent systematic efforts that have concluded that employing imputation in the analysis of single-cell RNA-seq data has no clear benefits as the zeroes observed in the data reflects both true gene expression and measurement error^53–55^, we chose to avoid an imputation step in our study. Of course, others may take different decisions and include imputation in preprocessing the input data in future studies. As this investigation is the first of its kind and we strived for simplicity at this stage, we have taken the measured gene expression values as is, but future studies may consider extending our approach considering the stochastic nature of the expression of different genes measured^57,58^, possibly preferring targets whose expression is more stable.

Even though the combinatorial optimization problems solved here are in the worst-case exponentially hard (NP-complete^33^ in computer science terminology), the actual instances that arise from the single-cell data could be either formally solved to optimality or shown to be infeasible with modern optimization software on the NIH Biowulf system, which has a hard limit of 240 wall-clock hours for any job. Of note, Delaney et al., have recently formalized a related optimization problem in analysis of single-cell clustered data for immunology^35^. Their optimization problem is also NP-complete in the worst case and they could solve sets of up to size four using heuristic methods^35^. We have shown that the optimal ILP solutions we obtained are often substantially smaller than solutions obtained via a greedy heuristic (**Figure 3**, **Supplementary Materials 8** including **Table S3**).

On the cautionary side, experiments with target gene sets that were further filtered by low expression in normal tissues showed that the individual target set problem can become infeasible in many instances. Even when the instance remained feasible, optimal cohort treatment set sizes increased rapidly as the expression levels allowed decreased (**Figure 6**), pointing to potential inherent limitations of applying such combination approaches to patients in the clinic and the need to carefully monitor their putative safety and toxicity in future applications. Finally, functional enrichment analysis of genes commonly occurring in the optimal target sets reinforced the central role of the widely studied oncogene *EGFR* and other transmembrane kinases. We also found that that the less-studied phosphatase *PTPRZ1* is a useful target, especially in brain cancer.

In summary, this study is the first to harness combinatorial optimization tools to analyze emerging single-cell data to portray the landscape of feasible personalized combinations in cancer medicine. Our findings uncover promising membranal targets for the development of future oncology medicines that may serve to optimize the treatment of cancer patient cohorts in several cancer types. The MadHitter approach and the accompanying software made public can be readily applied to address additional fundamentally related research questions and analyze additional cancer data sets as they become available.

## Methods

### Data Sets

We retrieved and organized data sets from NCBI’s Gene Expression Omnibus (GEO)^59^ and Ensembl’s ArrayExpess^60^ and the Broad Institute’s Single Cell Portal (https://portals.broadinstitute.org/single_cell). Nine data sets had sufficient tumor and non-tumor cells and were used in this study; an additional five data sets had sufficient tumor cells only and were used in testing early versions of MadHitter.. Suitable data sets were identified by searching scRNASeqDB^61^, CancerSea^62^, GEO, ArrayExpress, Google Scholar, the TISCH resource (http://tisch.comp-genomics.org) and the 10x Genomics list of publications (https://www.10xgenomics.com/resources/publications/). We required that each data set contain measurements of RNA expression on single cells from human primary solid tumors of at least two patients and the metadata are consistent with the primary data. We are grateful to several of the data depositing authors of data sets for resolving metadata inconsistencies by e-mail correspondence and by sending additional files not available at GEO or ArrayExpress.

We excluded blood cancers and data sets with single patients. The only exception is that we used one blood cancer data set (GSE132509) for a specific test of how we parameterized the killing thresholds. When it was easily possible to separate cancer cells from non-cancer cells of a similar type, we did so. In the specific case of GSE132509, there were two cell type annotations in GEO and a third annotation at (http://tisch.comp-genomics.org) and the malignant cells were far more consistent than the non-malignant cells. Therefore, for only the one analysis of GSE132509, we limited computations to the malignant cells that were consistent in the three annotations.

The main task in organizing each data set was to separate the cells from each sample or each patient into one or more single files. Representations of the expression as binary, as read counts, or as normalized quantities such as transcripts per million (TPM) were retained from the original data. When the data set included cell type assignments, we retained those to classify cells as “cancer” or “non-cancer”, except in the data set of Karaayvaz et al.^63^ where it was necessary to reapply filters described in the paper to exclude cells expressing few genes and to identify likely cancer and likely non-cancer cells. To achieve partial consistency in the genes included, we filtered data sets to include only those gene labels recognized as valid by the HUGO Gene Nomenclature Committee (http://genenames.org), but otherwise we retained whatever recognized genes that the data submitters chose to include. After filtering out the non-HUGO genes, but before reducing the set of genes to 1269 or 900 or 424 or 58, we filtered out cells as follows. Some data sets came with low expressing cells filtered out. To achieve some homogeneity, we filtered out any cells expressing fewer than 10% of all genes before we reduced the number of genes. As an exception, for the blood cancer data set (GSE132509) we found it necessary to raise the threshold to 20% because the data set had been noticeably less filtered than the main nine data sets. In **Supplementary Materials 4**, we tested the robustness of this 10% threshold. Finally, we retained either all available genes from among either our set of 1269 genes encoding cell-surface receptors that met additional criteria on low expression or available ligand-mimicking peptides.

### Sampling Process to Generate Replicates of Data Sets

As shown in **Table 1**, the number of cells available in the different single-cell data sets varies by three orders of magnitude; to enable us to compare the findings across different data sets and cancer types on more equal footing, we employed sampling from the larger sets to reduce this difference to one order of magnitude. This goes along with the data collection process in the real world as we might get measurements from different samples at different times. Suppose for a data set we have *n* genes, and *m* cells comprising tumor cells and non-tumor cells. We want to select a subset of *m′* < *m* cells. We select a set of *m′* cells uniformly at random without replacement from among all cells. Then we partition the selected cells into *m*_*t*_′ tumor cells and *m*_*n*_′ non-tumor cells to define one *replicate*. In most of the computational experiments shown we used 20 replicates and we report either the arithmetic mean or entire distribution of quantities such as the CTS size.

### Considering a previously defined set of target genes and of HPA gene expression across different normal tissues

The general aim of our methods is to target the cancer cells while sparing the adjacent non-cancer cells as much as possible. A related concern is that genes within the target set could be expressed at high levels in other normal tissues that are not part of the non-cancer cells from the tumor microenvironment included in the input data sets. One way to address this problem is to identify genes that have low expression in the majority of the tissues and to use them to obtain a target set. This approach has been pioneered in a recent paper on selecting gene targets suitable for CAR-T therapy^22^. The authors selected 533 candidate genes that they judged could be reasonable targets for CAR-T. They made this selection based on expression data from the Human Protein Atlas^76^ and the Genotype-Tissue Expression consortium (GTEx)^77^, which have expression information from multiple tissues which was used to identify low expressed target genes.

McKay et al.^22^ used a threshold of 15 TPM units of expression (written in their work as log2(TPM+1) ≤ 4), but they allowed a small number of tissues to exceed this threshold. Instead, we used quantitative levels of expression for finer granularity in analysis, as described in the next subsection. One clinical difference is that we looked only at adult tissues because we are analyzing adult tumors, while CAR-T therapy can be used for either childhood or adult tumors. The reason to focus on cell-surface receptors, as suggested by Dannenfelser et al.^21^, is that CAR-T therapy requires a cell-surface target that may or may not be a receptor, antibody technologies require a cell surface receptor, and the ligand-mimicking peptide nanotechnology that we summarized in the Introduction also requires cell surface receptor targets.

### Construction of target gene sets that are lowly expressed in normal tissues

To analyze the tissue specificity of the 1269 candidate target genes, the RNAseq based multiple tissue expression data was obtained from the Human Protein Atlas (HPA) database (https://www.proteinatlas.org/about/download; Date: May 29, 2020). The HPA database includes expression values (in units of transcripts per million (TPM)) for 37 tissues from HPA (rna_tissue_hpa.tsv.zip)^76^ and 36 tissues from the Genotype-Tissue Expression consortium (rna_tissue_gtex.tsv.zip)^77^. Next, to identify target genes with low or no expression within majority of adult human tissues, for the 1269 candidate genes we identified genes whose average expression across tissues is below certain threshold value (0.25, 0.5, 1, 2, 5, and 10 TPM) in both HPA and GTEx data sets. Using the intersection of low expression candidate genes from HPA and GTEx data sets, we generated lists of high confidence targets. The size of the resulting high confidence target genes varied from 424 (average expression less than 0.25 TPM) to 900 (average expression across tissue less than 10 TPM) genes (**Table 2**). While the total number of genes decreases slowly, the decrease is much steeper if one excludes olfactory receptors and taste receptors (**Table 2**). These sensory receptors are not typically considered as cancer targets, although a few of these receptors are selected in optimal target sets when there are few alternatives (**Figure 6**). MadHitter was run on all nine data sets using the expression information from the high confidence gene lists.

### Assembling Lists of Membrane Target Genes

We are interested in the set of genes *G* that i) have the encoded protein expressed on the cell surface and ii) for which some biochemistry lab has found a small peptide (i.e., amino acid sequences of 5-30 amino acids) that can attach itself to the target protein and get inside the cell carrying a tiny cargo of a toxic drug that will kill the cell and iii) encode proteins that are receptors. The third condition is needed because many proteins that reside on the cell surface are not receptors that can undergo RME. The first condition can be reliably tested using a recently published list of 2799 genes encoding human predicted cell surface proteins^76^; we reduced the list to 1269 by requiring that the proteins be receptors, which is necessary for RME-based therapies but not for CAR-T therpy^21^. For condition ii), we found two review articles in the chemistry literature^19–20^ that list targets effectively meeting this condition. Intersecting the lists meeting conditions i) and ii) gave us 38 genes/proteins that could be targeted (**Table 3**).

Most of the data sets listed in **Table 1** had expression data on 1200-1220 of these genes because the list of 1269 includes many olfactory receptor genes that may be omitted from standard genome-wide expression experiments. Among the 38 genes in **Table 3**, 13/14 data sets have all 38 genes, but GSE57872 was substantially filtered and has only 10/38 genes; since GSE57872 lacks non-tumor cells, we did not use this data set in any analyses shown.

Because the latter review^20^ was published in 2017, we expected that there are now additional genes for which ligand-mimicking peptides are known. We found 20 additional genes and those are listed in **Table 4**. Thus, our target set analyses restricted to genes with known ligand-mimicking peptides use 58 = 38 + 20 targets.

### Definition of the Minimum Hitting Set Problem and Solution Feasibility

One of Karp’s original NP-complete problems is called “hitting set” and is defined as follows^33^. Let *U* be a finite universal set of elements. Let *S*_1_, *S*_2_, *S*_3_, ..., *S*_*k*_ be subsets of *U*. Is there a small subset *H* ⊆ *U* such that for *i* = 1, 2, 3,...,*k*, *S*_*i*_ ∩ *H* is non-empty. In our setting, *U* is the set of target genes and the subsets *S*_*i*_ are the single cells. In reference 79, numerous applications for hitting set and the closely related problems of subset cover and dominating set are described; in addition, practical algorithms for hitting set are compared on real and synthetic data.

Among the applications of hitting set and closely related NP-complete problems in biology and biochemistry are stability analysis of metabolic networks^80–84^, identification of critical paths in gene signaling and regulatory networks^85–87^ and selection of a set of drugs to treat cell lines^88–89^ or single patients^90–91^. More information about related work can be found in **Supplementary Materials 1**.

Two different difficulties arising in problems such as hitting set are that 1) an instance may be *infeasible* meaning that there does not exist a solution satisfying all constraints and 2) an instance may be *intractable* meaning that in the time available, one cannot either i) prove that the instance is infeasible or ii) find an optimal feasible solution. *All instances of minimum hitting set that we considered were tractable* on the NIH Biowulf system. Many instances were provably infeasible; in almost all cases. we did not plot the infeasible parameter combinations. However, in **Figure 4**, the instance for the melanoma data set with the more stringent parameters was infeasible because of only one patient sample, so we omitted that patient for both parameter settings in **Figure 4**.

### Basic Optimal Target Set Formulation

Given a collection *S* = {*S*_1_,*S*_2_, *S*_3_,...} of subsets of a set *U*, the hitting set problem is to find the smallest subset *H* ⊆ *U* that intersects every set in *S*. The hitting set problem is equivalent to the set cover problem and hence is NP-complete. The following ILP formulates this target set problem:

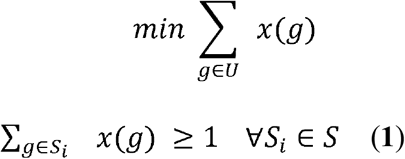

In this formulation, there is a binary variable *x*(*g*) for each element *g* ∈ *U* that denotes whether the element *g* is selected or not. Constraint (**1)** makes sure that from each set *S*_*i*_ in S, at least one element is selected.

For any data set of tumor cells, we begin with the model that we specify a set of genes that can be targeted, and that is *U*. Each cell is represented by the subset of genes in *U* whose expression is greater than zero. In biological terms, a cell is killed (hit) if it expresses at any level on one of the genes that is selected to be a target (i.e., in the optimal target set) in the treatment. In this initial formulation, all tumor cells are combined as if they come from one patient because we model that the treatment goal is to kill (hit) all tumor cells (all subsets). In a later subsection, we consider a fair version of this problem, taking into account that each patient is part of a cohort. Before that, we model the oncologist’s intuition that we want to target genes that are overexpressed in the tumor.

### Combining Data on Tumor Cells and Non-Tumor Cells

To make the hitting set formulation more realistic, we would likely model that a cell (set) is killed (hit) only if one of its targets is overexpressed compared to typical expression in non-cancer cells. Such modeling can be applied in the nine single-cell data sets that have data on non-cancer cells to reflect the principle that we would like the treatment to kill the tumor cells and spare the non-tumor cells.

Let *NT* be the set of non-tumor cells. For each gene *g*, define its average expression *E*(*g*) as the arithmetic mean among all the non-zero values of the expression level of *g* and cells in *NT*. The zeroes are ignored because many of these likely represent dropouts in the expression measurement. Following the design of experiments in the lab of N. A., we define an expression ratio threshold factor *r* as a real number whose baseline value is 2.0. We adjust the formulation of the previous subsection, so that the set representing a cell (in the tumor cell set) contains only those genes *g* such that the expression of *g* is greater than *r* × *E*(*g*) instead of greater than zero. We keep the objective function limited to the tumor cells, but we also store a set to represent each non-tumor cell, and we tabulate which non-tumor cells (sets) would be killed (hit) because for at least one of the genes in the optimal target set, the expression of that gene in that non-tumor cell exceeds the threshold *r* × *E*(*g*). We add two continuous parameters *lb* and *ub* each in the range [0,1] and representing respectively a lower bound on the proportion of tumor cells killed and an upper bound on the proportion of non-tumor cells killed. The parameters *lb, ub* are used only in two constraints, and we do not favor optimal solutions that kill more tumor cells or fewer non-tumor cells, so long as the solutions obey the constraints.

More explicitly, if we let *E*(*g, C*) be the expression level of gene *g* in cell *C*, then *g* is in the set of genes that can cover *C* if and only if *E*(*g, C*) ≥ *r* × *E*(*g*). We inspect this relationship for every pair of genes and cells to come up with the hitting set instance of our ILPs.

Our choice of the lower bound (*lb*) on the proportion of tumor cells killed in the range [0.7,0.9] is somewhat arbitrary at this stage, but it is motivated by the widely used response evaluation criteria in solid tumors (RECIST) revised guidelines version 1.1^92^. Those criteria include the category of partial response (PR), which is usually classified together with complete response (CR) as a responder when one makes a dichotomy between response and non-response. PR is defined as “At least a 30% decrease in the sum of diameters of target lesions, taking as reference the baseline sum diameters.” Because volume is proportional to the cube of diameter, a 30% decrease in the diameter sum corresponds to a decrease of at least 66% in the volume, which we round up to 70% (0.7). Because the tumor mass contains both tumor cells and non-tumor cells and we want to kill the tumor cells disproportionately, we increased the baseline threshold for *lb* to 80% (0.8) and considered values as high as 0.95 or even 0.99 in some supplementary analyses. In any case, *lb* is a user-controlled parameter and can be set as the user of our software deems appropriate. Since most cancer treatments have some toxicity, we set the baseline value of *ub* to the low nonzero value 0.1; in some analyses, we use the lower value 0.05. If one wanted to distinguish the different types of non-tumor cells, one could add different upper bound parameters for each cell type.

### The Fair Cohort Target Set Problem for a Multi-Patient Cohort

We want to formulate an integer linear program that selects a set of genes *S*\* from available genes in such a way that, for each patient, there exists an individual target set 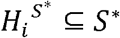 of a relatively small size (compared to the optimal ITS of that patient alone which is denoted by *H*(*i*)).

Let *U = {g*_1_, *g*_2_, ..., *g*_*|U|*_} be the set of genes, where |*U*| denotes the number of elements in the universe of genes *U*. There are n patients. For the *i*^*th*^ patient, we denote by *S*_*P(i)*_, the set of tumor cells related to patient *i*. For each tumor cell *C* ∈ *S*_*P(i)*_, we describe it as a set of genes which is known to be targetable to cell *C*. That is, *g* ∈ *C* if and only if a drug containing *g* can target the cell *C*. In the ILP, there is a binary variable *x*(*g*) corresponding to each gene *g* ∈ *U* that shows whether the gene *g* is selected or not. There is a binary variable *x*(*g, P(i)*) which shows whether a gene *g* is selected in the target set of patient *P*(*i*). Let *α* be an adjustable positive integer, which we interpret as a slack for each patient’s solution. The objective function is to minimize the total number of genes selected, subject to having a target set of size at most *H*(*i*) + *α* for patient *P*(*i*) where 1 ≤ *i* ≤ *n* (constraint (2)).

Constraint (3) ensures that, for patient *P*(*i*),we do not select any gene *g* that are not selected in the global set.

Constraint (4) ensures all the sets corresponding to tumor cells of patient *P*(*i*) are hit.

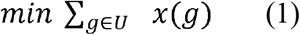

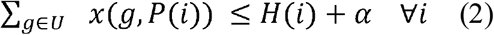

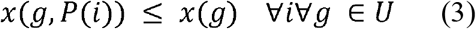

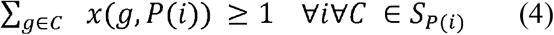

In this ILP, there is a variable corresponding to each gene. Additionally, there is a variable per each pair of gene and patient, therefore, the total number of variables is |***U***|(***n + 1***). The total number of constraints in the form of constraints (2), (3), and (4) are ***n, n***|***U***|, and 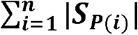 respectively.

### Parameterization of the Fair Cohort Target Set Problem

In the Fair Cohort Target Set ILP shown above, we give more preference towards minimizing number of genes needed in the CTS. However, we do not take into account the number of non-tumor cells killed. Killing (covering) too many non-tumor cells potentially hurts patients. In order to avoid that, we add an additional constraint to both the ILP for the local instances and the global instance. Intuitively, for patient *P*(*i*), given an upper bound of the portion of the non-tumor cell killed *ub*, we want to find the smallest cohort target set *H*(*i*) with the following properties:

1. *H*(*i*) covers all the tumor cells of patient *P*(*i*).
2. *H*(*i*) covers at most *ub* * |*NT*_*P(i)*_| where *NT*_*P(i)*_ is the set of non-tumor cells known for patient *P*(*i*); the binary variable *y*(*C*) represents whether the cell C is covered.

The ILP can be formulated as follows:

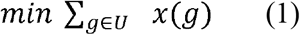

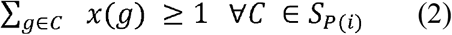

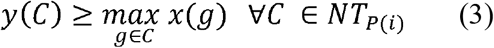

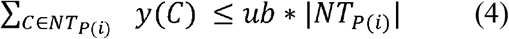

In the abovementioned ILP, the total number of variables is |*U*| + |*S*_*P(i)*_ + *NT*_*P(i)*_|, where there is a variable corresponding to each gene and a variable corresponding to each cell of patient *P*(*i*). The total numbers of constraints in the form of constraints (2), (3), and (4) are |*S*_*P(i)*_|, |*NT*_*P(i)*_|. |*U*|, and 1 respectively. In the actual ILP implementation, constraint (3) is in fact |*U*| different constraints for each *C* ∈ *NT*_*P(i)*_.

With this formulation, the existence of a feasible solution is not guaranteed. However, covering all tumor cells might not always be necessary either as we discussed above in the context of RECIST criteria for tumor response to treatment. Hence, we add another parameter *lb* to let us model this scenario. In the high-level, this is the ratio of the tumor cells we want to cover. The ILP can be formulated as follows:

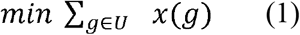

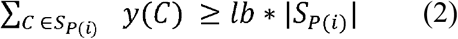

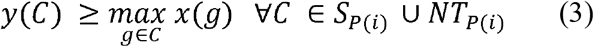

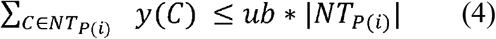

Notice that constraint (2) here is different from the one above as we only care about the total number of tumor cells covered. In the above ILP, the total number of variables is |*U*| + |*S*_*P(i)*_| + |*NT*_*P(i)*_|, where there is a variable corresponding to each gene and a variable corresponding to each cell of patient *P*(*i*). The total numbers of constraints in the form of constraints (2), (3), and (4) are 1, (|*S*_*P(i)*_| + |*NT*_*P(i)*_|). |*U*|, and 1 respectively.

Even with both *ub* and *lb*, the feasibility of the ILP is still not guaranteed. However, modeling the ILP in this way allows us to parameterize the ILP for various other scenarios of interest. While the two ILPs above are designed for one patient, one can extend these ILPs for multi-patient cohort. In a way that is similar to how we define *x*(*g, P(i)*) the binary variable *y*(*C, P(i)*) denotes whether the cell *C* is covered for the i^th^ patient.

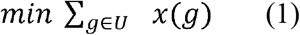

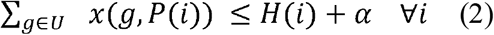

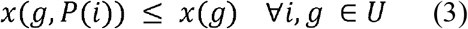

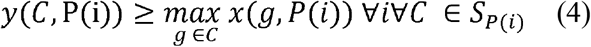

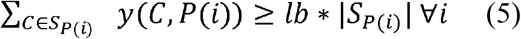

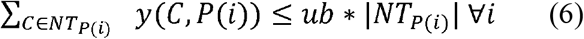

In the abovementioned ILP, the total number of variables is 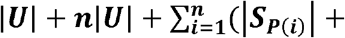 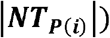 where there is a variable corresponding to each gene, a variable per each pair of gene and patient, and a variable corresponding to each cell of each patient ***P(i)***. The total numbers of constraints in the form of constraints (2), (3), (4), (5), and (6) are 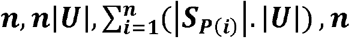, ***n***, and ***n*** respectively.

### Implementation Note, Accounting for Multiple Optima and Software Availability

We implemented in Python 3 the above fair cohort target set formulations, with the expression ratio r as an option when non-tumor cells are available. The parameters *α, lb, ub* can be set by the user in the command line. To solve the ILPs to optimality we usually used the SCIP library and its Python interface^93^. To obtain multiple optimal solutions of equal size we used the Gurobi library (https://www.gurobi.com) and its python interface. When evaluating multiple optima, for all feasible instances, we sampled 50 optimal solutions that may or may not be distinct, using the Gurobi function select_solution(). To determine how often each gene or pair of genes occur in optimal solutions, we computed the arithmetic mean of gene frequencies and gene pair frequencies over all sampled optimal solutions.

The software package is called MadHitter. The main program is called hitting_set.py. We include in MadHitter a separate program to sample cells and generate replicates, called sample_columns.py. So long as one seeks only single optimal solutions for each instance, exactly one of SCIP and Gurobi is sufficient to use MadHitter. We verified that SCIP and Gurobi give optimal solutions of the same size. If one wants to sample multiple optima, this can be done only with the Gurobi library. The choice between SCIP and Gurobi and the number of optima to sample are controlled by command-line parameters use_gurobi and num_sol, respectively. The MadHitter software is available on GitHub at https://github.com/ruppinlab/madhitter

## Abbreviations

CTS: cohort target set, synonym of global hitting set
GEO: Gene Expression Omnibus
GHS: global hitting set, synonym of cohort target set
GTEx: Genotype-Tissue Expression (project or consortium)
HPA: Human Protein Atlas
HUGO: Human Genome Organization
IHS: individual hitting set, synonym of individual target set
ILP: integer linear programming
ITS: individual target set
lb: lower bound on fraction of tumor cells killed
RME: receptor-mediated endocytosis
TPM: transcripts per million
ub: upper bound on fraction of non-tumor cells killed

## Acknowledgements

This research is supported in part by the Intramural Research program of the National Institutes of Health, National Cancer Institute. This research is supported in part by the University of Maryland Year of Data Science Program. This research is supported in part by start-up funds from Northwestern University and a research award from Amazon to support the research of S.K. This work utilized the computational resources of the NIH HPC Biowulf cluster. (http://hpc.nih.gov). Thanks to E. Michael Gertz for technical assistance with SCIP, Gurobi, and Biowulf. Thanks to Allon Wagner, Keren Yizhak and Sushant Patkar for assistance in identifying and retrieving suitable single-cell RNAseq data sets. Thanks to Leandro Hermida for technical advice.

## Competing Interests

The authors declare that they have no competing interests.

## Notes

https://github.com/ruppinlab/madhitter

### Competing Interest Statement

The authors have declared no competing interest.

### Summary of Updates

Modified title. Substantially rewritten Abstract. New results on targets of CAR-T therapy. New background information to justify the modeling choices. A little bit of redistribution of content between the main document and the supplementary document.

