## Supplementary Information for "The Landscape of Receptor-Mediated Precision Cancer Combination Therapy: A Single-Cell Perspective"

### Supplementary Materials

#### 1. Related Work in Combinatorial Optimization and Superselectivity

Combining the paradigms of personalized medicine, modular treatment strategies, and single-cell data sets, we formulate a two-layered combinatorial optimization problem that models a modular approach to personalized cancer treatment. We seek to make principled estimates of quantities such as: How many different treatments are needed to kill all cells in a tumor? If one can kill all cells in a tumor, what fraction of normal cells nearby will also get killed as a side effect of the combination therapy?

In this subsection, we review two types of related work in the area of combinatorial optimization. The first type of related work is applications of a problem called “hitting set” in bioinformatics. The second type of related work is other multi-layered problems in combinatorial optimization (defined and described below). At the end, we also review some research on physical chemistry and nanotechnology that helps justify how we model the expression ratio $r$.

One of Karp’s original NP-complete problems is called “hitting set” and is defined as follows^1^. Let $U$ be a finite universal set of elements. Let $S_{1}, S_{2}, S_{3}, ..., S_{k}$ be subsets of $U$**.** Is there a small subset $H \subseteq U$ such that for $I=1, 2, 3, ..., k$**,** $S_{i}\cap H$ is non-empty. In reference 2, numerous applications for hitting set and the closely related problems of subset cover and dominating set are described; in addition, practical algorithms for hitting set are compared on real and synthetic data. Among the applications of hitting set and closely related NP-complete problems in biology and biochemistry are stability analysis of metabolic networks^3-7^, identification of critical paths in gene signaling and regulatory networks^8-10^ and selection of a set of drugs to treat cell lines^11,12^ or single patients^13,14^.

The hitting set problem has a dual problem called “set cover” or “subset cover”, which is also among Karp’s list of original NP-complete problems. In the subset cover problem, the input again consists of a universe $U$ and a set of subsets $S_{1}\subset U, S_{2}\subset U, ...$ possibly with weights. The goal is to find a minimum cardinality or minimum weighted collection of subsets whose union is $U$. The set cover problem and variations of it with additional constraints or objective function terms has been used extensively in bioinformatics. Several prior uses have been in cancer studies^15-21^. Two of these set cover-based methods, called UNCOVER^20^ and NETPHIX^21^ have some resemblance with our formulation of the CTS problem. These methods assume that the data are patients or cell lines with known somatic or germline mutations, rather than our assumption that the input is gene expression data. Given mutation data on patients and knowledge of which drugs are effective for which gene mutations, UNCOVER and NETPHIX can then use integer linear programming (ILP) techniques to solve a variant of set cover that finds optimal drug combinations for the entire set of patients, but treatments are not optimized for individual patients as in our ITS formulation. NETPHIX adds information about gene and protein networks to favor gene sets that are functionally related. An example of set cover in the study of non-cancer complex diseases can be found in reference 22.

The NP-completeness of the hitting set and subset cover problems implies that there is unlikely to be a polynomial-time algorithm that finds the optimal solution to arbitrary instances. However, several of the studies cited in the previous paragraphs and our study also show that large real instances of hitting set and subset cover can be solved to optimality via techniques from ILP. Figure S1 shows a toy example hinting at why finding optimum-size solutions of the hitting set instances we formulate may be non-trivial.

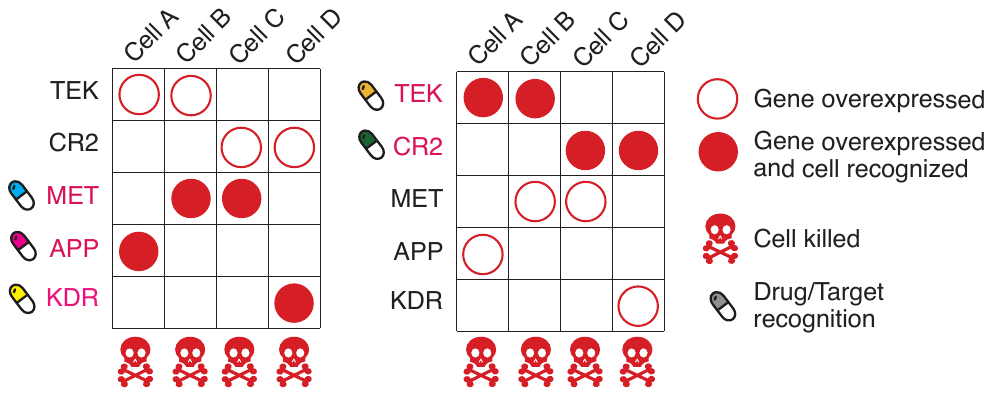

Figure S1. A schematic small example of killing a four tumor cells illustrating why choosing a minimum-size combination of targets may be non-trivial. The schematic tumor has four cancer cells (A, B, C, D in separate columns), which may express any of five cell-surface receptor genes (rows) that may be targeted selectively by modular treatments (pills). If one targets {APP, KDR, MET}, all cancer cells will be killed (left panel). However, if, instead, one would target {CR2, TEK} then all cancer cells in the given example tumor will be killed with just two targets (right panel) instead of three, providing a smaller solution.

We review briefly the four papers on using hitting set to select drugs since these are most pertinent to our work. In the initial formulation, the set $U$ is a set of drugs, the subsets $S_{i}$ represent cell lines; a drug $d$ is a member of $S_{i}$ if the cell line responds (typically by being less viable) to drug $d$^11^. Thus, the underlying data (edges) for the bipartite graph representation of hitting set are binary response data for each possible pair of (cell line, drug). The biological objective is to find a minimum number of drugs such that each cell line responds to at least one drug^11^.

Mellor et al. pointed out that in practice the response to a drug is incomplete and the cell line may evolve to resist the drugs, so one would really like to solve a t-hitting set problem in which is required that $|S_{i} \cap H| \geq t \geq1$, for a fixed $t$, rather than $>0$^12^. Moreover, they pointed out that for the cell-line drug-response data used in reference 11, the t-hitting set problem is fixed-parameter tractable and used this observation to find hitting sets of size 3 in a few seconds of computer time^12^.

Vera-Licona et al. and Pang et al. considered the more clinical problem of treating actual patients rather than cell lines^13,14^. Generally, they modeled the potential drug targets as a set of genes that were labeled as either “on-target” (meaning that a drug targeting such a gene would ameliorate the disease) or “off target” (meaning that a drug targeting such a gene would cause side effects). Via mixed integer linear programming (MILP) they solved various optimization problems that ensure that all source-sink paths in a network are covered by drugs targeting on-target-genes while seeking to minimize the number of off-target genes that the selected drugs would affect^13,14^. Importantly for what follows, each problem instance they considered was for a single network of genes representing either one patient or one genomically homogeneous cohort of patients.

In this work, we introduce a new aspect of global optimization across a cohort of patients. Developing the target-specific part of each immunotoxin ligand, nanoparticle, or degrader is expensive, so we would prefer to reduce the size of $U$**.** On the other hand, we would like to find an optimal or near-optimal hitting set for the single cells of each patient. And that leads to a two-layer optimization problem.

Given a set of instances $I_{1}, I_{2}, I_{3}, ...$ of hitting set problems that have the same universal set *U***,** find a set of optimal solutions $H_{1},H_{2},H_{3},...$such that the union of $H_{1},H_{2},H_{3},...$ is as small as possible. That union of solutions conceptually represents the set of targeted drugs that one would need to develop to ensure than an optimal treatment plan was available for each patient. We call this problem “Fair Hitting Set (FHS)” based on the intuition that we want a high-quality solution for all patients that does not require one patient to receive many more nanoparticles than necessary for that patient, while other patients receive optimal treatments.

There are previous theoretical studies on other combinatorial optimization problems with an analogous two-layer structure. The best known of these is the “Universal Traveling Salesman Problem (TSP)”^23-25^ in which one is given $n$ points with distances defined in a metric space (typically the Euclidean plane) and one seeks a cyclic ordering (called a “tour”) of all $n$ points such that the implied subtour for any subset of $k$ points is not too much worse than then optimal tour for those $k$ points. Jia and colleagues extended this type of study to another network analysis problem called Steiner tree^23^. Marcolino and colleagues applied these methods to another network problem called “influencing a social network” when the structure of the network is unknown^26^.

From the perspective of algorithms, the work that is closest to ours is also in reference 23 that studied the problem of set cover, which is the dual of hitting set. In set cover, as mentioned above, the universe $U$ again consists of $n$ elements and an instance is a set $S=\{S_{1},S_{2},...\}$ of subsets of $U$. One seeks the smallest collection of $S_{i}$ who union is $U$. Jia and colleagues showed asymptotically matching upper and lower bounds for universal set cover that the worst-case best solution grows as $n^{1/2}$ times the cost of the optimal solution for each subset of $U$. They also extended their results to a weighted version of hitting set. Instead of the worst-case formulation, Grandoni et al. and Adamczyk et al. proposed two alternative probabilistic formulations of universal set cover in which one tries to optimize the expected performance on the worst subset^27,28^. Based on prior work, it would have been natural to use the name “universal hitting set” for the problem we study, but this name has already been used by others for a combinatorial optimization problem in biological sequence analysis^29^.

One of the most interdisciplinary aspects of our work is how we use progress in the theory and experiments of superslectivity^30^ which comes from the fields of physical chemistry and nanotechnology to model how future cancer treatments can recognize overexpressed cell surface receptors, as modeled by the parameter $r$. Validated nanoparticle systems^31^ use multivalency of the ligand-mimicking peptides and the principle of superselectivity to increase the probability that at least one peptide will bind to the target. Martinez-Veracochea and Frenkel^30^ did theoretical calculations and molecular simulations to suggest that the relationship looks like $b=cv^{ɤ}+o(v^{ɤ})$, where $b$ is the number of bound surface protein molecules, $c$ is some tunable “constant”, $v$ is the number of ligand-mimicking peptides also called the “valency” and ɤ is some exponent > 1 that is tunable to some extent. What we write here as the exponent ɤ they called α, but we use the symbol α for the fairness parameter. The higher the achievable value of ɤ, the more superselective the system is because a small change in the valency ($v$) leads to a much larger change in the number of bound protein molecules. Another way to measure the effect of multivalency is the ratio of the probability of binding of a multivalent particle to the probability of binding to a monovalent particle; this ratio is denoted by β^32^. In a cell-free system, Dubacheva and colleagues^31^ validated the exponential behavior in physical experiments and showed that ɤ in the range of 2 to 3 is achievable. In a different cell-free system, using binding to DNA, Estirado and colleagues^32^ demonstrated superselectivity, with β as high as 10^5^.

One biomedical non-cancer setting in which superselectivity is potentially important is when viruses bind host cell receptors. For the important case of influenza A virus binding to glycans, Overeem and colleagues recently demonstrated in vitro that superselectivity occurs and visualized values of ɤ in the range of 2 to 10 depending on the receptor glycan density^33^.

Superselectivity in the context of nanomedicine was reviewed recently by Woythe and colleagues^34^. To date, no nanoparticles with multivalent binding have been approved. For our purposes, we are mostly interested in proteins at a cell surface being the binding targets. Binding to nucleic acid targets such as microbial genomes has also been studied and has also been shown to have the phenomenon of superselectivity^32,35^.

Superselectivity has been modeled based on methods from statistical physics. In this terminology, it is the increased “permutation entropy” created by the multivalent nanoparticles that leads to increased probability of binding between nanoparticles and receptors^34^. The strength of binding affects the probability that binding will lead to endocytosis; weak and flexible binding peptides are best for this purpose^34^. Theoretical analysis shows that the plot of probability of binding (y-axis) as a two-variable function of i) receptor density (i.e., protein expression) and ii) number of ligands per nanoparticle approaches a step function and that the inflection point is adjustable. For example, in several of the curves in Figure 3 of reference 34 a doubling of the receptor density changes the probability of binding from < 0.1 to 1. These curves justify to some extent our default value of $r=2$ for the proportion of gene target overexpression that leads to binding between a decorated nanoparticle and the protein target on the cancer cell surface.

By requiring some force at an angle normal (i.e., perpendicular) to the target surface to release the content of the nanoparticle, one can move from “superselectivity” to “hyperselectivity” in which the binding function acts almost like an on-off switch^34,36^. The effect of the force is to increase the exponent ɤ from [2,3] to approximately 6^36^. By tuning the amount of force, one can get very close to a Heaviside step function, which represents the idealized behavior of an on-off switch^36^. The prediction of hyperselectivity remains theoretical and wet lab tests are needed to validate and quantify the tuning methods suggested^36^. If validated, then any $r>1$ should be achievable, but it is natural that values of r corresponding to the jumps in the polynomial superselective curves would be tested first.

The above summary considers nanoparticles with multivalent ligand-mimicking peptides as decoration to deliver the toxic drugs. For the delivery of CAR-T, the issue of selectivity is beginning to be considered. Using combinatorial logic, Hernandez-Lopez and colleagues demonstrated in vitro feasibility of r in the range 5-10 with a behavior that resembles at least superselectivity if not hyperselectivity^37^. These studies of nanoparticle systems and CAR-T justify our decisions to test primarily $r$in the range 1.5-3.0 (Section 3) and then to extend the range up to 5.0 (Section 7).

#### 2. Robustness Analysis of CTS Size as a Function of Number of Cells Sampled

In this subsection, we show CTS size as a function of the number of cells sampled in eight data sets partitioned into four data sets each in Figures S2 and S3. For the four data sets in Figure S2, the levels start to plateau at 500 cells and above and that is one reason we selected these data sets for most of the analyses shown in the main document. For the four data sets illustrated in Figure S3, 250 cells are usually sufficient. For the remaining data set (brain cancer (GSE89567)), which is not in Figures S2 and S3, the data were highly filtered by the submitters, the CTS size is close to 1 with all parameter values we tried and hence 250 cells are sufficient.

We also show that the mean ITS sizes and mean CTS sizes are not substantially different when using all cells as compared to sampling (Table S1). We prefer the sampling approach as it makes the different data sets more comparable and it shows that for future data sets, sampling 500 cells should be sufficient to capture the heterogeneity of gene expressions across single cells.

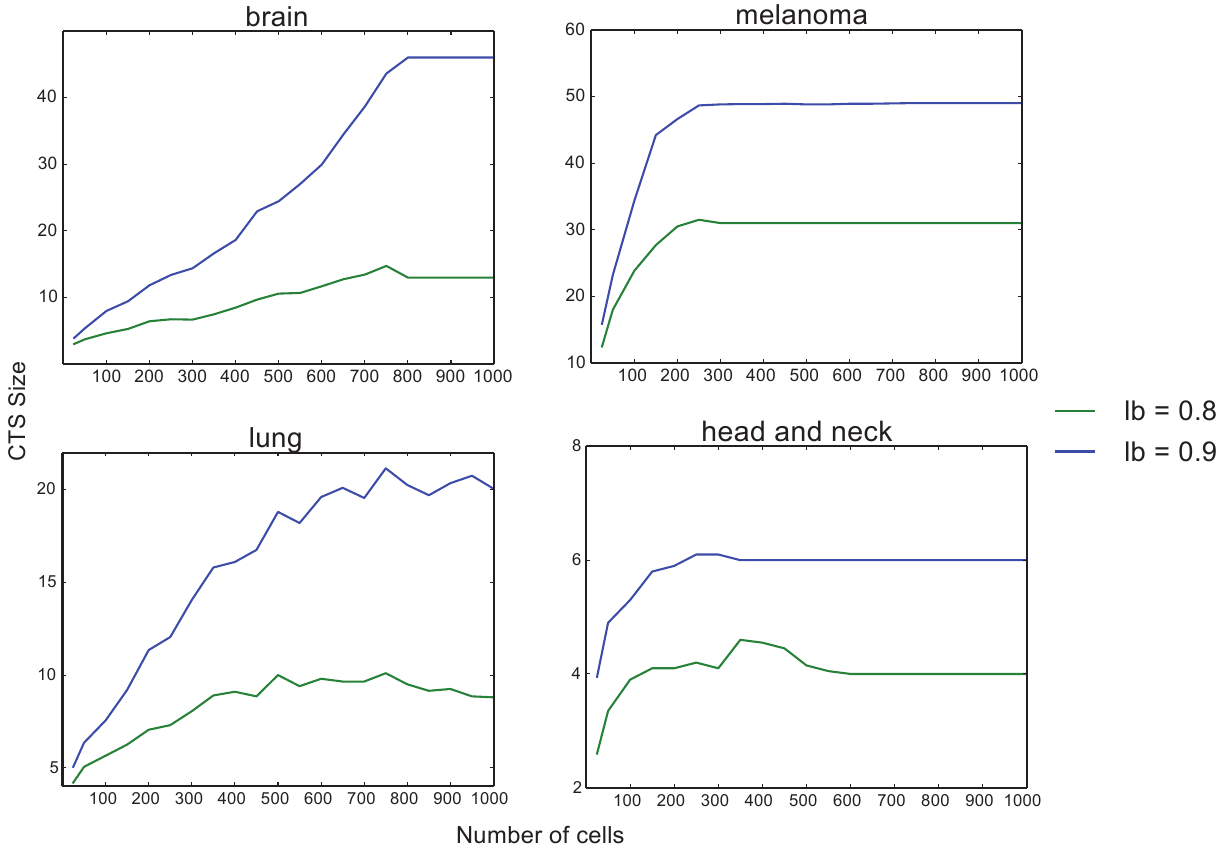

Figure S2. Mean CTS sizes in 20 replicates of the brain (GSE84465), melanoma, lung, and head and neck cancer data sets as a function of number of cells sampled and using baseline parameters.

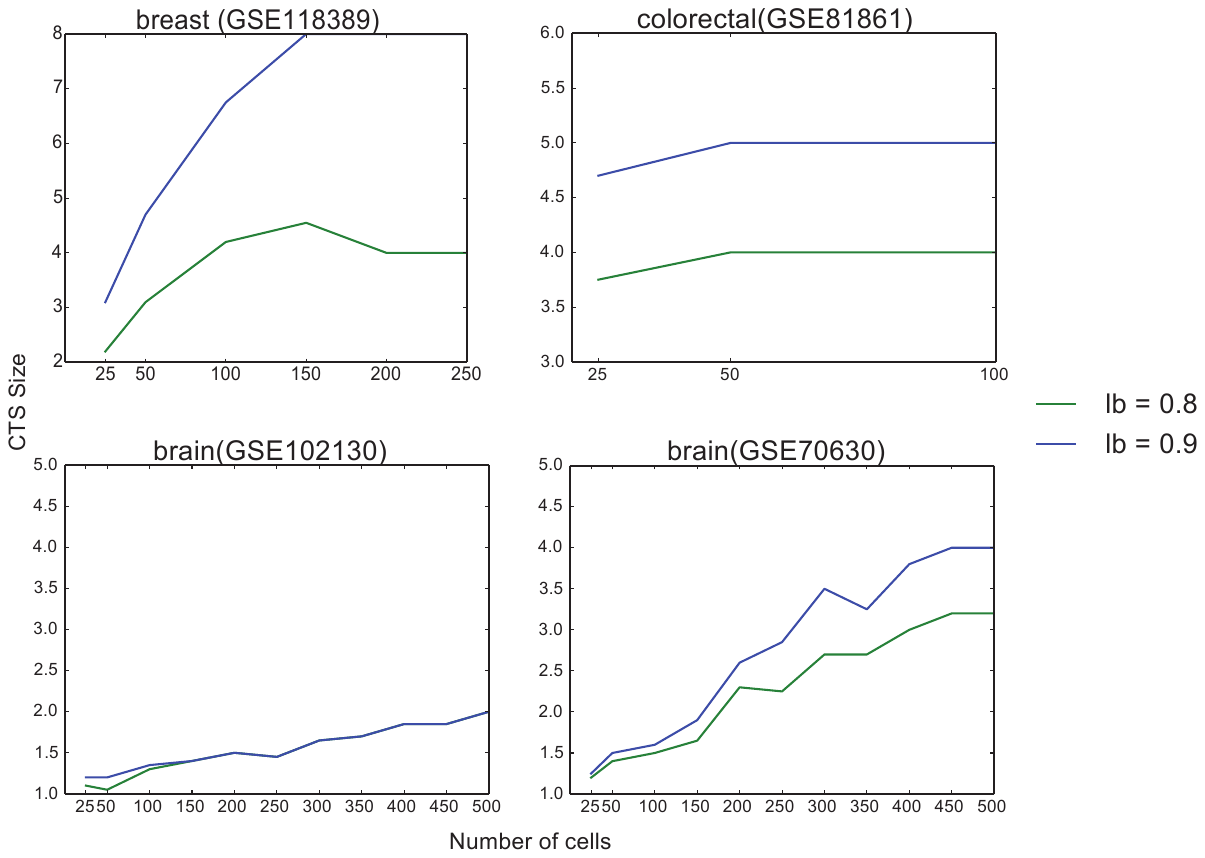

Figure S3. Mean CTS sizes in 20 replicates of the breast, colorectal, and brain (GSE102130 and GSE70630) data sets as a function of number of cells sampled and using baseline parameters.

Table S1. Comparison of cohort target set (CTS) sizes and individual target set (ITS) set sizes based on sampling a fixed number of cells or using all cells. The sampling mean CTS size is the mean over 20 replicates. The sampling mean ITS size is the mean over 20 replicates and all patients. The all cells mean ITS size is the mean over all patients. Even though each replicate in our sampling protocol considers only a subset of cells, the CTS or ITS for that replicate can be larger than for all cells because of randomness.

| Tumor Type | Data Set | Sampling Mean CTS | All Cells CTS | Sampling Mean ITS | All Cells Mean ITS |
| --- | --- | --- | --- | --- | --- |
| Breast | GSE118389 | 4 | 4 | 2.67 | 2.67 |
| Brain | GSE89567 | 1 | 1 | 1 | 1 |
| Brain | GSE70630 | 3.2 | 3 | 1.233 | 1.33 |
| Brain | GSE102130 | 2 | 2 | 1 | 1 |
| Brain | GSE84465 | 10.6 | 13 | 3.9125 | 4.75 |
| Colorectal | GSE81861 | 5 | 5 | 2.33 | 1.833 |
| Head and neck | GSE103322 | 4.45 | 4 | 1 | 1.077 |
| Melanoma | GSE115978 | 31 | 31 | 2.67 | 2.409 |
| Lung | E-MTAB-6149 | 10 | 9 | 2.98 | 3 |

#### 3. The Landscape of CTS Size for Additional Data Sets

In the main text, we showed heatmaps of CTS size as a function of $lb$, the lower bound on proportion of tumor cells killed and of $ub$, the upper bound on the proportion of non-tumor cells killed for four data sets. Here, in Figures S4-S8, we show analogous plots for five other data sets. These data sets tend to have few patients, fewer cells and/or more aggressive filtering of cells by the data submitters. Most of these five data sets do not show a sharp increase in CTS size as $lb$ increases or as $ub$ decreases.

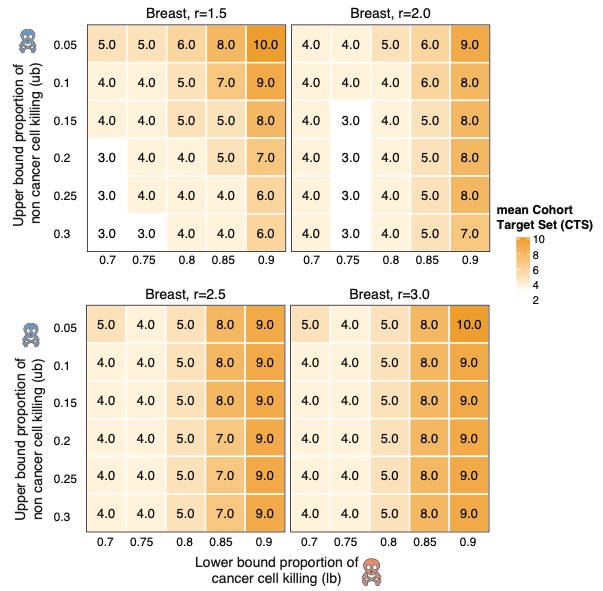

Figure S4. Heat maps of mean CTS size for a breast cancer data set (GSE118389). Plotted values are in the range [3,10].

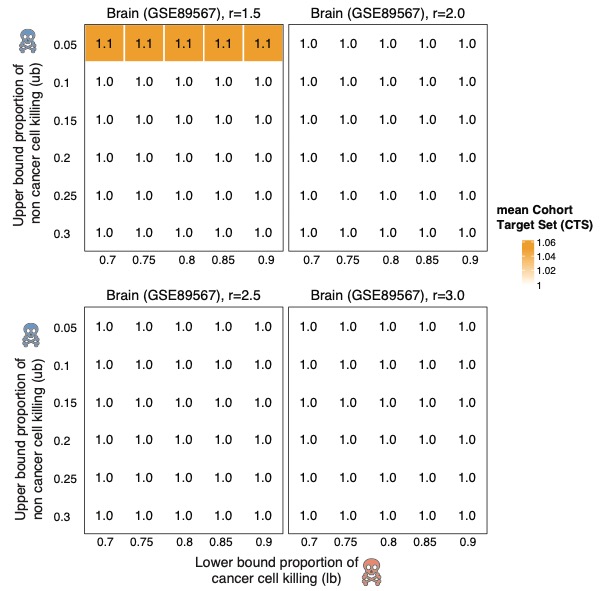

Figure S5. Heat maps of mean CTS size for a second brain cancer data set (GSE89567). Plotted values are in the range [1, 1.1]$.$

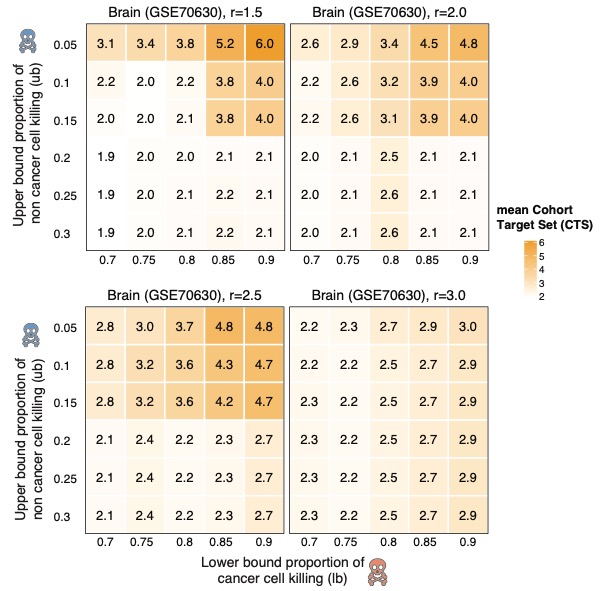

Figure S6. Heat maps of mean CTS size for a third brain cancer data set (GSE70630). Plotted values are in the range [1.85, 6].

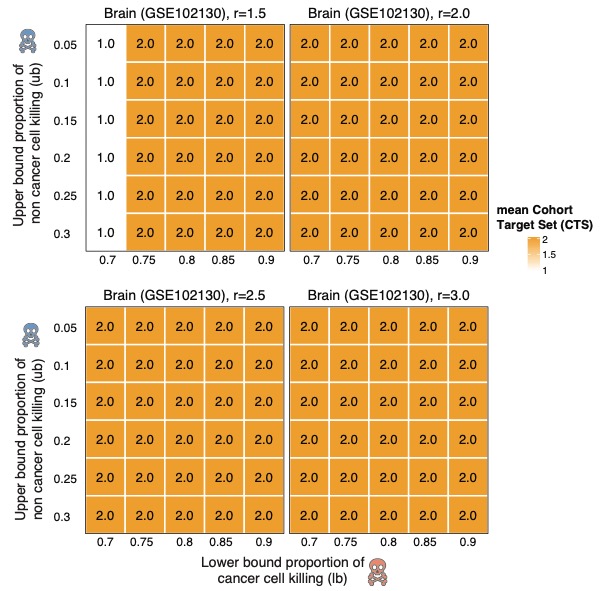

Figure S7. Heat maps of mean CTS values for a fourth brain cancer data set (GSE102130). Plotted values are in the range [1, 2.0], but vary only when the expression ratio $r=1.5.$

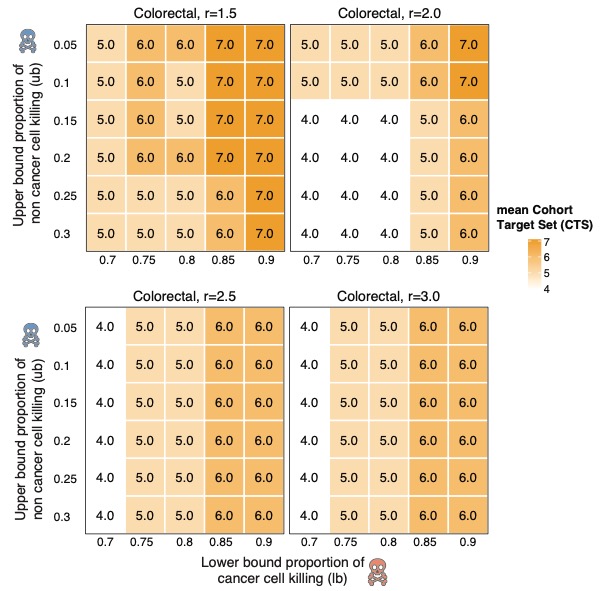

Figure S8. Heat maps for mean CTS values of a colorectal cancer data set (GSE81861). Plotted values are in the range [4,7].

#### 4. The Qualitative Transitions in Cohort Set Sizes Arise Regardless of the Threshold for Filtering Low-Expressing Cells

For the analyses of the main text, we tried to homogenize the data sets to some extent by filtering out cells that express fewer than 10% of all genes at some level. For at least two of the data sets (brain cancer and lung cancer) analyzed in Figure 2 and the breast cancer data set, we observed a sharp increase in CTS size as $lb > 0.8$ or $ub < 0.1$. We considered the possibility that the observed phased transitions are an artifact of the filtering being insufficiently stringent. The analyses in Figures S9-S11 increased the filtering threshold to 15%, 20%, 25% for tumor cells in the brain, lung, and breast cancer data sets. These supplementary analyses show that the phase transitions persist. Not surprisingly, the CTS sizes decrease markedly when more low-expressing cells are removed from the input data and higher-expressing cells that are easier to target are retained.

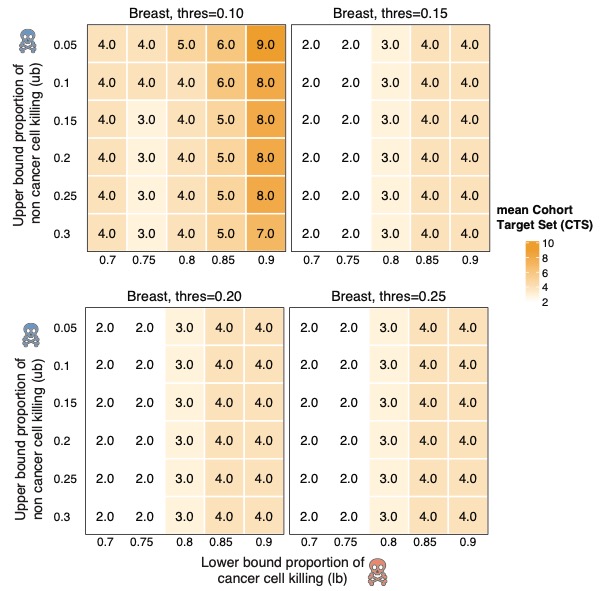

Figure S9. CTS sizes for a breast cancer data set (GSE118389). The different panels represent different stringencies of filtering out tumor cells that express few genes. The filtering threshold is above each panel.

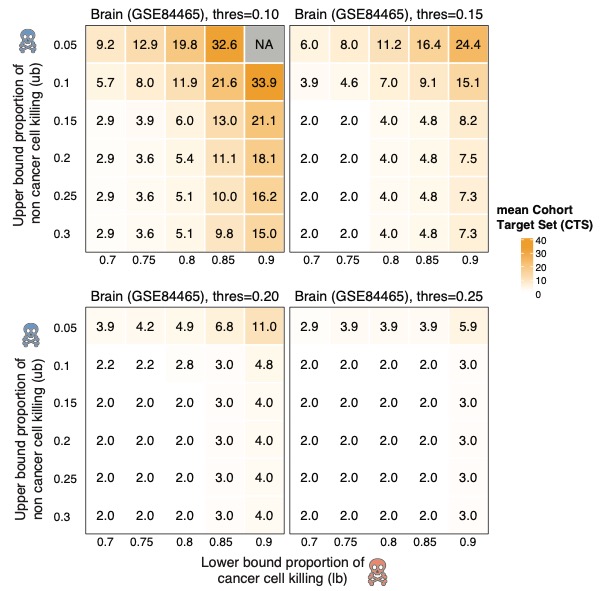

Figure S10. CTS sizes for the same brain cancer data set (GSE84465) analyzed in Figure 2 and elsewhere in the main text. The different panels represent different stringencies of filtering out tumor cells that express few genes. The filtering threshold is above each panel.

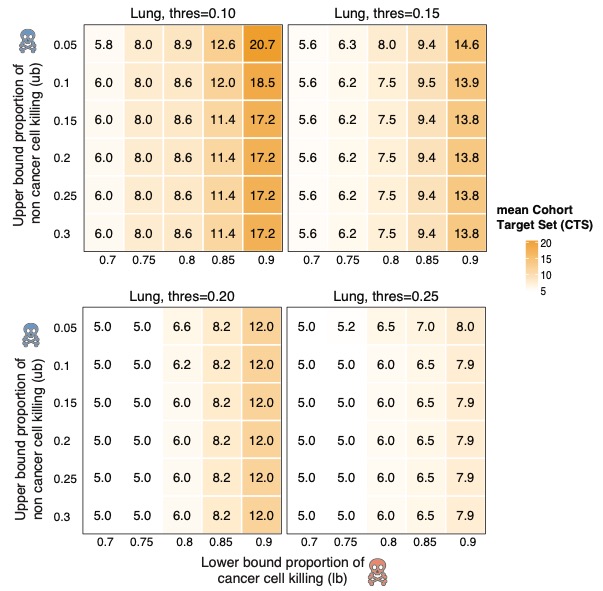

Figure S11. CTS sizes for the same lung cancer data set (E-MTAB-6149) analyzed in Figure 2 and elsewhere in the main text. The different panels represent different stringencies of filtering out tumor cells that express few genes. The filtering threshold is above each panel.

#### 5. Distributions of ITS Sizes for Additional Data Sets

Here, we show distributions of ITS sizes for the data sets not included in the main Figure 3 and for feasible extreme setting of the cell killing stringency thresholds. Description of the greedy algorithm and more results can be found later in Supplementary Section 8. In general, we observe that the ITS size is similar for the ILP and greedy algorithms when it is small, but as the instances get harder, the MadHitter ILP can sometimes find optimal solutions that are much smaller than the greedy algorithm. A small amount of the variation in ITS sizes is due to random sampling across the 20 replicates for each set of parameter values.

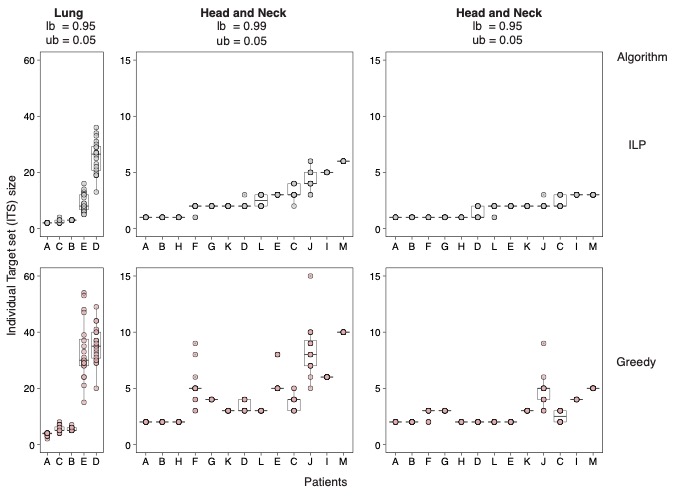

Figure S12. Plots of patient-specific ITS sizes for the optimal ILP and the greedy algorithm for two additional data sets in which the instances remain feasible for extreme thresholds (*lb* = 0.095 or 0.99, *ub*= 0.05, and *r*= 2), but which were omitted from the main Figure 3. For each patient, we sampled 20 replicates as described in Methods.

#### 6. Heatmaps of CTS Size for More Stringent Cell Killing Thresholds

In the main text and Supplementary Materials 3, we showed heatmaps of CTS size for $lb$varying from 0.7 up to 0.9 and for $ub$ varying from 0.3 to 0.05. In this subsection, we show as Figures S13-S14 heatmaps of CTS size for more extreme values of $lb$ up to 0.99 and for eight out of nine data sets. For these plots, we keep the expression ratio $r$ fixed at the baseline value of 2.0.
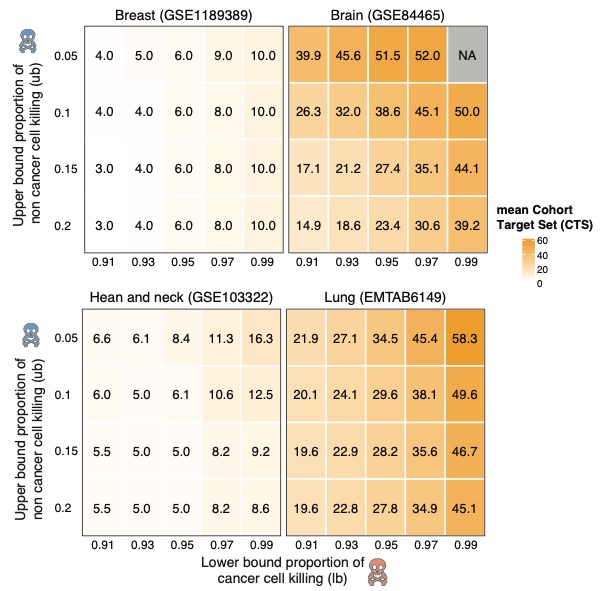
Figure S13. CTS sizes for four data sets and more stringent killing thresholds than in main Figure 2 and Figures S4-S8. Here, $lb$, the lower bound on the fraction of tumor cells targeted ranges from 0.91 to 0.99. NA stands for Not Available because there is no feasible solution.

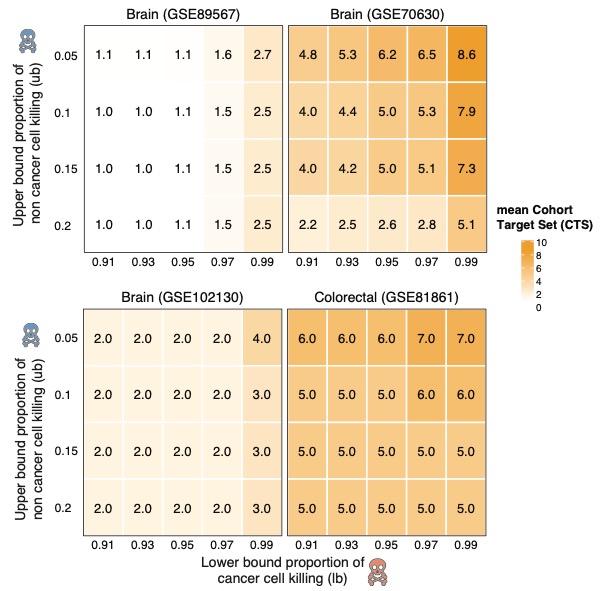

Figure S14. CTS sizes for four data sets different from those in Figure S13 and with more stringent killing thresholds as in Figure S13.

#### 7. CTS Size as a Function of the Expression Ratio $\boldsymbol{r}$

In the main text and earlier subsections of Supplementary Materials, we described and plotted how CTS size varies as a function of $lb$and $ub$, while mostly keeping the expression ratio $r$ fixed. In this subsection, we vary $r$in increments of 0.25 while keeping $lb = 0.8$and $ub = 0.1$fixed at baseline values. Figures S15-S16 plot in two dimensions how CTS size varies. Table S2 lists for each data set, the tested value of $r$that yields the lowest value of CTS size. In the future, it would be preferable to select $r$in a gene-specific manner^38^ taking into account target-specific modeling and experiments about the affinity for the target-specific part of the treatment (e.g., ligand-mimicking peptide or antibody) for the target receptor^31-33^. The rationale for considering values of $r$ up to 5 is explained at the end of the first subsection of the Supplementary Material.

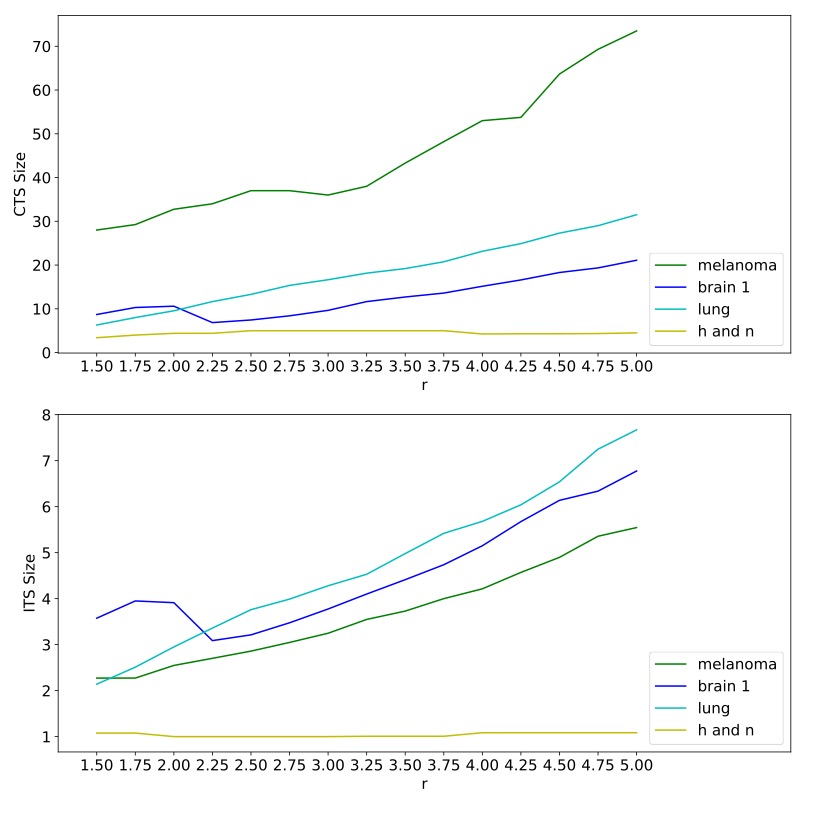

Figure S15. Mean CTS sizes and ITS sizes with $ub=0.1, lb=0.8$ and $r$ varying in the range [1.5,5.0] for the four data sets analyzed in Figure S2 and most of the main text: brain (GSE84465), head and neck (GSE103322), melanoma (GSE115978) and lung (E-MTAB-6149).

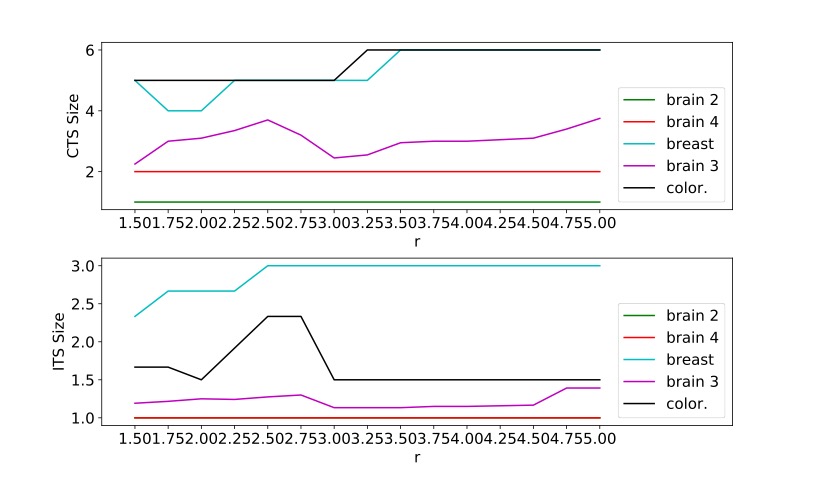

Figure S16. Mean CTS sizes and ITS sizes with $ub=0.1, lb=0.8$ and $r$ varying in the range [1.5,5.0] for the five datasets analyzed in Figures S4-S8: breast (GSE118389), brain 2 (GSE89567), brain 3 (GSE70630), brain 4 (GSE102130), and colorectal (GSE81861).

Table S2. We varied the expression ratio $r$ from 1.5 to 5.0 in increments of 0.25, while keeping other parameters at baseline values. We report one value or a range of consecutive values that give the lowest CTS size. The CTS size is the mean of 20 replicates of sampled cells and hence may have a non-integer value.

| Tumor Type | Data Set | Best $r$ | CTS Size at best $r$ |
| --- | --- | --- | --- |
| breast | GSE118389 | 1.75-2.0 | 4.00 |
| brain | GSE89567 | 1.50-5.0 | 1.00 |
| brain | GSE70630 | 1.50 | 2.25 |
| brain | GSE102130 | 1.50-5.00 | 2.00 |
| brain | GSE84465 | 2.25 | 6.85 |
| colorectal | GSE81861 | 1.50-3.0 | 5.00 |
| head and neck | GSE103322 | 1.50 | 3.40 |
| melanoma | GSE115978 | 1.50 | 28.0 |
| lung | E-MTAB-6149 | 1.50 | 6.30 |

#### 8. A Greedy Algorithm for Selecting Target Gene Sets as an Alternative Benchmark

An ILP solver can take substantial time; SCIP and Gurobi do take substantial time on some of our HS instances. Therefore, we implemented another algorithm that requires little time, but might still give us good solutions in some cases. Since the decisions made by the algorithm come from a simple way of ranking the genes, we call this algorithm a *greedy algorithm* (see detailed description below).

We ran the greedy algorithm against all nine data sets. If it had turned out that the results from our greedy algorithm, in terms of CTS size had been as good as, or close to the results from ILP solver, then the merit in using resources to solve ILP would be mainly to have confidence that the solutions we obtained are optimal. In most data sets, the CTS from the greedy algorithm are much larger than the minimum-size CTS produced by the ILP solver (Table S3). Hence, the effort to develop the ILP method is justified in practice.

Table S3. Comparison of the cohort and individual target set sizes output from ILP (optimal) and greedy algorithm, among 1269 targets, $lb= 0.8, ub=0.10, r = 2.$Individual target set sizes are much larger with the greedy algorithm than with the optimal ILP algorithm. Cohort target set sizes are larger for eight out of nine data sets (exception GSE103322) even though the greedy algorithm considers the entire cohort and could trade off a higher ITS size for a lower CTS size. As explained in conjunction with Figures S2-S3, we selected for each data set a round number of cells to sample that strikes a balance between how many cells are available and where the CTS size begins to plateau.

| Tumor Type | Data Set | Cells sampled | ILP CTS Size | Greedy CTS Size | ILP ITS Size | Greedy ITS Size |
| --- | --- | --- | --- | --- | --- | --- |
| Breast | GSE118389 | 250 | 4 | 7 | 2.67 | 3.33 |
| Brain | GSE89567 | 250 | 1 | 2.05 | 1 | 1.95 |
| Brain | GSE70630 | 500 | 3.2 | 5.8 | 1.233 | 2.442 |
| Brain | GSE102130 | 500 | 2 | 2.0 | 1 | 1.667 |
| Brain | GSE84465 | 500 | 10.6 | 22.7 | 3.9125 | 7.188 |
| Colorectal | GSE81861 | 100 | 5 | 6 | 2.33 | 2.67 |
| Head and neck | GSE103322 | 500 | 4.45 | 2.05 | 1 | 1.923 |
| Melanoma | GSE115978 | 500 | 31 | 59.6 | 2.473 | 5.67 |
| Lung | E-MTAB-6149 | 500 | 10 | 17.55 | 2.98 | 5.7 |

Now we describe our greedy algorithm implementation more formally. We first describe the implementation for a single patient. We then show how to extend the implementation for a cohort of multiple patients.

Let $\Gamma=\{g_{1},g_{2},...,g_{|\Gamma|}\}$ be the (arbitrarily) ordered set of genes. Also, define ${\Gamma^{2}}\subseteq\Gamma\times\Gamma$ be the ordered set of pairs of genes. For each pair $p=(g_{i}, g_{j\geq i})$, we compute the set of tumor cells $T_{p}$ and the set of non-tumor cells $NT_{p}$ hit by $p$. We sort pairs in $\Gamma^{2}$ by the numbers of tumor cells hit by them (in decreasing order), breaking ties by the numbers of non-tumor cells targeted by them (in increasing order). If two pairs are equal, on both primary and secondary sorting keys, then we break ties using the order we have in $\Gamma$. The rationale for considering pairs rather than single genes is that there exist high-throughput robotic systems to evaluate pars of drug treatments simultaneously and we expected that considering two genes at a time would be more effective than one gene at a time. The first (smallest) pair in the ordered set ${\Gamma^{2}}$ is probably the most useful one as it should kill the highest number of tumor cells.

After we have the ordered set ${\Gamma^{2},}$we will construct the solution set $HS$ as follows. We will consider pairs, one by one, starting from the highest-ranking pair to the lowest-ranking pair. When we consider $p=(g_{i}, g_{j\geq i})$, we will consider adding $g_{i}, g_{j}$ to $HS$, in respective order. When considering the gene $g_{,}$ we will add $g$ to $HS$ if a) $g$hits at least one cell not hit by the previously selected genes and b) the cardinality of non-tumor cells killed by $HS\cup\{g\}$ does not exceed the upper bound, defined by $ub*|NT|$. We stop the process as soon as $HS$ kills at least $lb*|T|$ tumor cells or as soon as we run out of genes to add. If we run out of genes, then the greedy algorithm fails to find a feasible solution, but this case did not arise in the nine datasets considered.

We can take different states of $HS$ at different steps in our algorithm as partial solutions (of size bounded by $k$ for any constant $k$). This gives us information that we cannot get by running the ILP solver.

When we have a cohort of patients, it is not enough to consider the number of tumor cells killed and the number of non-tumor cells killed (or the summation of them). One potential problem arises when we have a difference in magnitude of available data from different patients. For example, a gene target that kills 10 out of 10 tumor cells of patient A and 500 out of 1000 tumor cells of patient B might be better than another gene target that kills 5 out of 10 tumor cells of patient A and 550 out of 1000 tumor cells of patient B. Instead, we use what we call *average effectiveness* as a criterion. We define effectiveness of a target for a patient as the ratio of tumor cells killed to the number of tumor cells. Average effectiveness of a target gene is an arithmetic mean of effectiveness of that target gene across all patients. In the example above, the average effectiveness of the first target is 0.75, while it is 0.525 for the second target. By using average effectiveness as the key for sorting, we obtain the unified order of ${\Gamma^{2}}$.

We then run the same algorithm for each patient separately, using the same gene pairs in the same order. The final output of our algorithm is the union of the targets selected for each patient.

#### 9. Comparisons of CTS Sizes for 1269 Cell Surface Targets and 58 Targets with Known Peptide-Mimicking Ligands and Analysis with 57 Targets of CAR-T Therapy

In this subsection, we show heatmaps (Figures S17 -19) in which the outcome measure is the difference CTS Size (1269 targets) - CTS Size (58 targets). We also list the 57 genes encoding targets of CAR-T therapy. As explained in the main text, differences can be negative when the smaller set of 58 forces targets to be reused by multiple patients. ITS Size (58 targets) must be greater than or equal to ITS Size (1,269 targets).

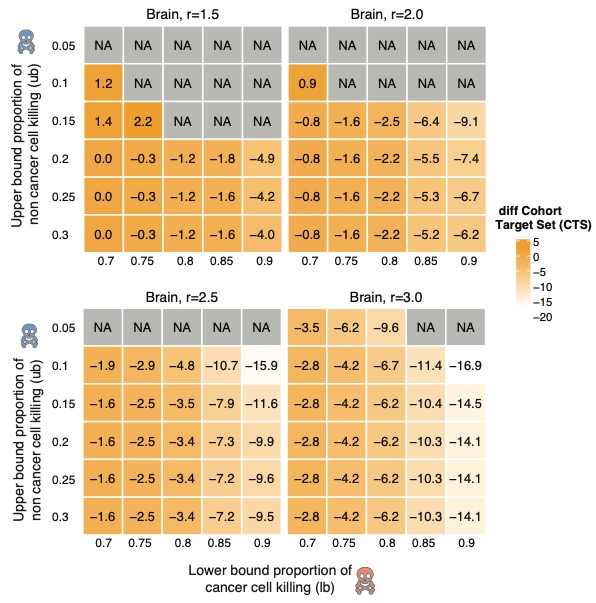
Figure S17. Difference of mean CTS sizes using 1269 target genes and 58 target genes for a brain cancer data set GSE84465. Some instances are infeasible (NA) for only 58 targets.

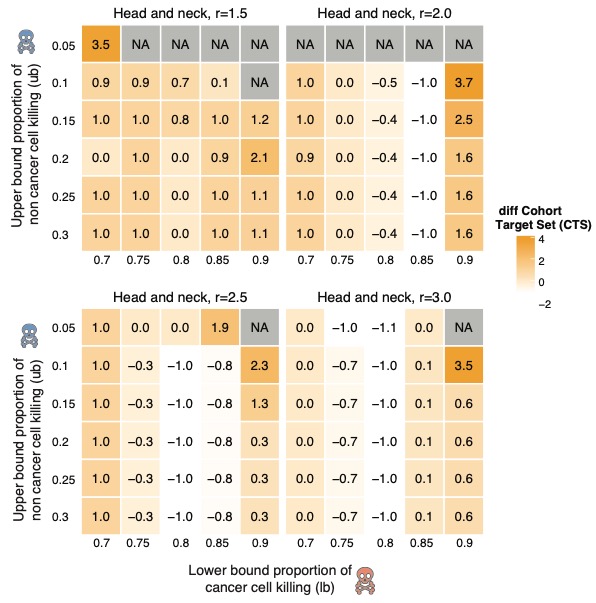

Figure S18. Difference of mean CTS sizes using 1269 target genes and 58 target genes for a head and neck cancer data set GSE103322. Some instances are infeasible (NA) for only 58 targets.

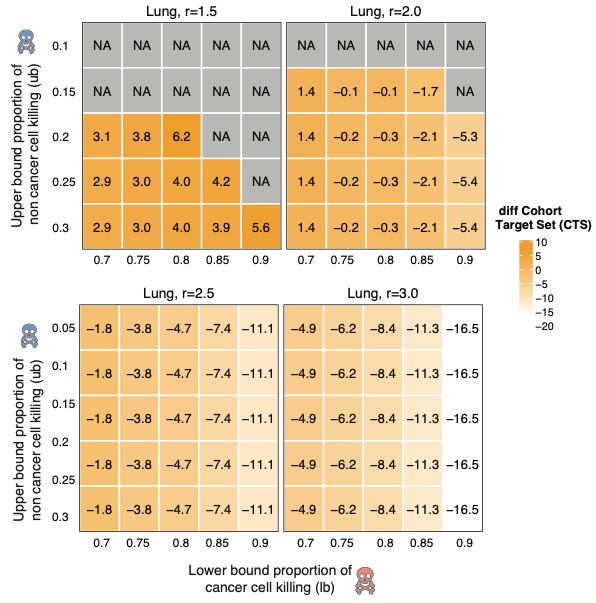

Figure S19. Difference of mean CTS sizes using 1269 target genes and 58 target genes for a lung data set E-MTAB-6149. Some instances are infeasible for only 58 targets.

#### Figure 2 in reference 39 lists in protein or lipid form 64 cell surface targets being tested in trial of CAR-T therapy. In Supplementary Table S4, we list 57 genes encoding the single protein targets, ignoring protein complexes and cell surface lipids. As described briefly in the main text, we used these 57 genes as an alternative input gene list to MadHitter.

Table S4. Single genes encoding single proteins that are being tested as targets of CAR-T therapy^39^. If the protein target is being tested in any of the six solid tumor types for which we analyzed a single cell data, then those tumor types are listed in the rightmost column.

| Protein(s) | Gene Symbol | Relevant Tumor Type(s) |
| --- | --- | --- |
| CD19 | *CD19* |  |
| BCMA | *BCMA* |  |
| CD22 | *CD22* |  |
| CD20 | *MS4A1* |  |
| CD123 | *IL3RA* |  |
| CD30 | *TNFRSF8* |  |
| CD38 | *CD38* |  |
| CD33 | *CD33* |  |
| CD138 | *SDC1* |  |
| CD56 | *NCAM1* |  |
| CD7 | *CD7* |  |
| CLL-1 | *CLEC12A* |  |
| CD10 | *MME* |  |
| CD34 | *CD34* |  |
| CS1 | *SLAMF7* |  |
| CD16 | *FCGR3A* |  |
| CD4 | *CD4* |  |
| CD5 | *CD5* |  |
| IL-1-RAP | *IL1RAP* |  |
| ITGB7 | *ITGB7* |  |
| TACI | *TNFRSF13B* |  |
| TRBC1 | *TRBC1* |  |
| MUC1 | *MUC1* | brain, lung, colon, breast |
| NKG2D | *NKG2D* | colon, breast |
| PD-L1 | *CD274* | brain, lung, colon, breast |
| CD133 | *PROM1* | brain, lung, colon, breast |
| CD117 | *KIT* |  |
| CD70 | *CD70* | breast, melanoma |
| ROR1 | *ROR1* | lung, breast |
| AFP | *AFP* |  |
| AXL | *AXL* |  |
| CD80 | *CD80* | lung |
| CD86 | *CD86* | lung |
| DLL3 | *DLL3* | lung |
| DR5 | *TNFRSF10B* |  |
| FAP | *FAP* |  |
| FBP, FRa | *FOLR1* |  |
| LMP1 | *PDLIM7* |  |
| MAGE-A1 | *MAGEA1* | lung |
| MAGE-A4 | *MAGEA4* | lung |
| MG7 | *PTGS2* |  |
| MUC16 | *MUC16* |  |
| PMEL | *PMEL* | melanoma |
| ROR2 | *ROR2* |  |
| VEGFR2 | *KDR* | melanoma |
| CD171 | *L1CAM* |  |
| CLD18 | *CLDN18* |  |
| EPHA2 | *EPHA2* | brain |
| EGFR and isoforms | *EGFR* | brain, liver, lung, colon, head and neck |
| PSCA | *PSCA* | lung |
| cMET | *MET* | colon, breast, melanoma |
| IL13Ra2 | *IL13RA2* | brain |
| EPCAM | *EPCAM* | colon, breast |
| PSMA | *FOLH1* |  |
| GPC3 | *GPC3* | lung |
| HER2 | *ERBB2* | brain, lung, colon, breast, head and neck |
| Mesothelin | *MSLN* | lung, breast |

Among the 57 genes, 23 were consistently present across all nine data sets. Using our default parameters and the mean over 20 replicates, we estimated what proportion of cancer cells would be killed if each individual target were used (Figure S20). Among the 23 genes, only *EGFR, AXL,* *MET, IL1RAP,* and *PROM1* might have high enough expression in any of the six tumor types we studied to have any reasonable chance of succeeding clinically, Several of the 23 genes are being considered in tumor types we did not analyze.

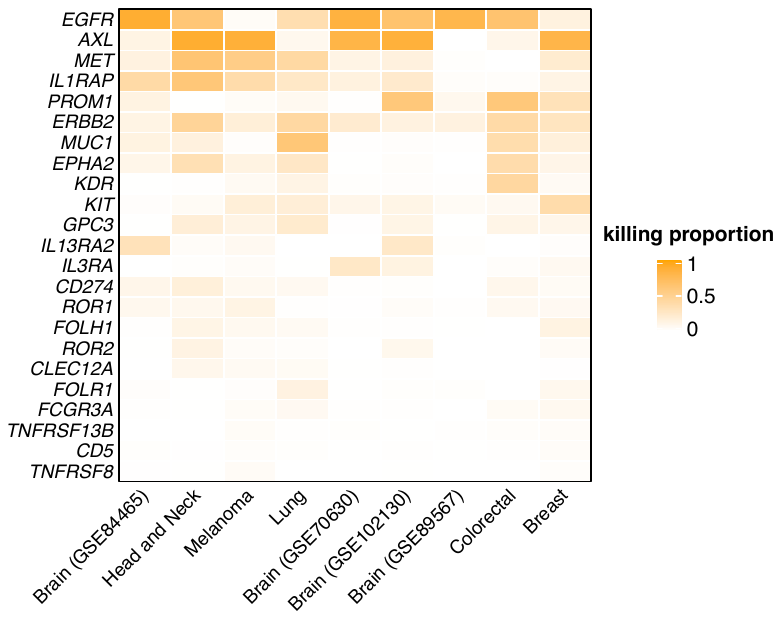

Figure S20. The proportion of cancer cells killed across the nine datasets by each of the 23 cell surface targets being tested in trial of CAR-T therapy.

#### 10. Additional Empirical and Theoretical Analysis of Fair Cohort Sets Including Divergence of Empirical Behavior and Theoretical Worst-Case Behavior

Figures S21-S25 show how the sizes of CTS and ITS change as a function of $\alpha(given baseline parameters)$. Recall that α is the number of extra treatments that a patient may receive above the minimum number. The analysis of the melanoma data set (Figure 5C) shows that the cohort target size can decrease substantially by allowing $\alpha$ to be greater than 0, while the average number of treatments that each individual patient receives varies little and is in the single digits. In general, we are interested in the smallest value of α at which the CTS size reaches its minimum in each column of the heatmaps and we see that this is always ≤ 4. The number of treatments per patient should be viewed as an upper bound because in our ILP formulation, we treat $\alpha$ as a constraint on all patients. One could add a tie-breaking term to the objective function to minimize the sum of the number of treatments given to all patients, but this would make the ILP harder to solve to optimality.

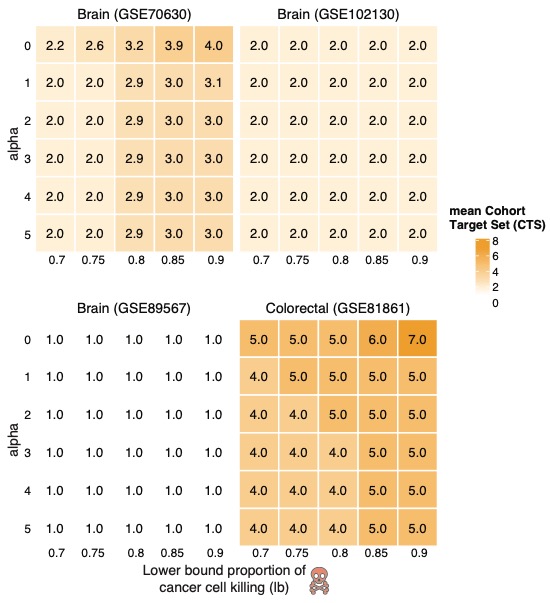

Figure S21. For four data sets not shown in Figure 6C, CTS sizes as a function of $\alpha, lb$.

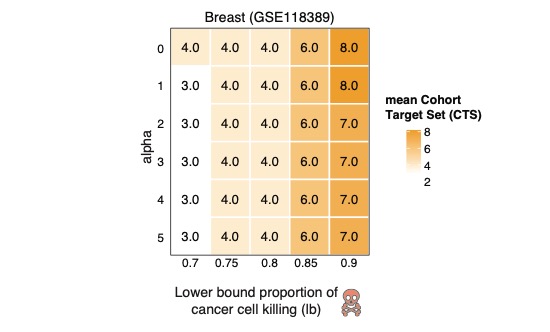

Figure S22. CTS sizes as a function of α, *lb* for the ninth data set.

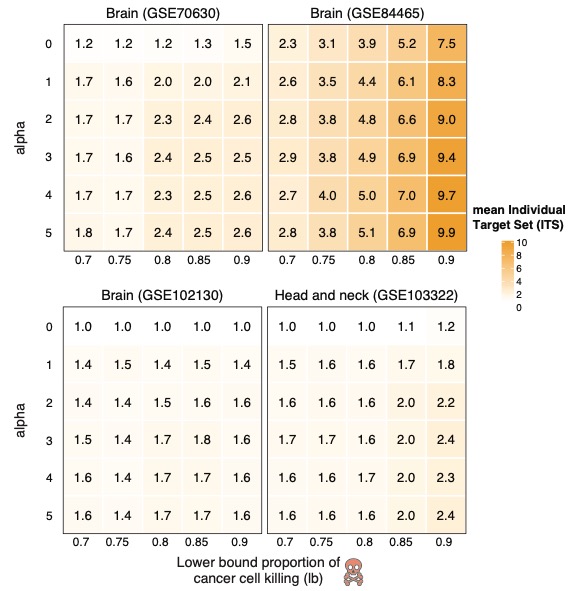

Figure S23. Mean ITS sizes as a function of α, *lb*.

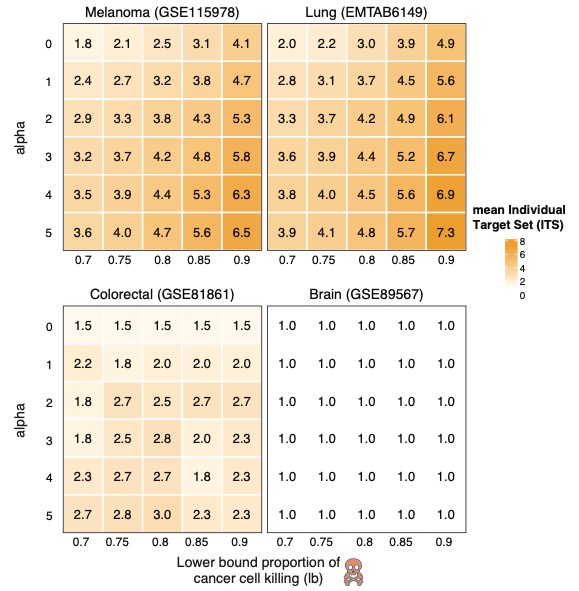

Figure S24. Mean ITS sizes as a function of α, *lb* for four additional data sets.

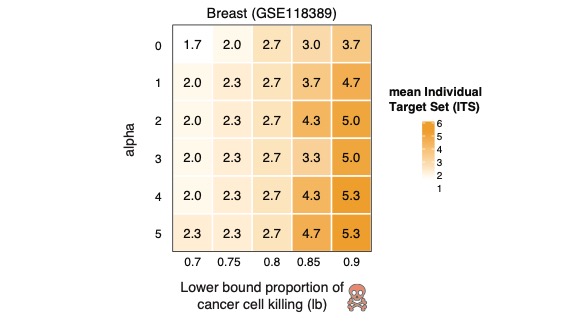

Figure S25. Mean ITS sizes as a function of α, *lb* for the ninth data set.

In contrast to the empirical data, a theoretical worst-case analysis matches the intuition that “each patient is different.” We give an example to show that in the worst-case instance, it is impossible to have a hitting set that is small and fair. Precisely, we construct an instance as follows. Suppose we have n patients and m tumor cells. Let $S$ be the set of tumor cells. The set of cells for every patient is identical. For $1\leq i\leq n$, let $e_{i}$be a unique gene that can kill all tumor cells for the i^th^ patient. For $j \in S$, let $r_{j}$ be a gene that can kill the tumor cell $j$ for every patient. The small optimal cohort target set size has size $m$ while the fair solution has size $n$. We can set $m=\sqrt{n}$ and we would need $\alpha$to be at least $\sqrt{n}-1$to get a cohort target set of size less than $n$.

**11.** **Optimal Solutions with Varying Upper Bounds on Gene Expression in Healthy Tissues**

To target genes whose expression is low in normal tissues to avoid collateral damage, we restricted the targetable genes to those with expression below certain thresholds (0.25, 0.5, 1, 2, 5, and 10 transcripts per million (TPM)) in healthy tissue (Methods). This restriction results in reduction of the size of targetable genes (Table 2). In Figure S29, we show that reduction of the size of targetable genes makes it difficult for MadHitter to find a feasible solution; the trend is different in different datasets and for different lower bounds. All runs summarized in the section were done with default settings of $r=2.0, ub=0.10, lb=0.80.$In the main Figure 6, the 6A and 6B panels are heatmaps which show the genes and their frequency of occurrence in the optimal solutions across different expression threshold. Figures S26-S28 represent the same type of heatmap in three additional datasets. Independent of the genes, the size of the optimal solution increases with decrease in size of targetable genes which is shown at a cohort level in Figure S30A, and at an individual level in Figures S30B and S31.

We also ran MadHitter on 533 target genes suggested for CAR-T therapy by MacKay et al.^39^. These genes are lowly expressed in most but not all tissues using a threshold of TPM=15. The 533 genes do not necessarily encode receptors. All instances in all nine data sets are feasible with the restricted 533-gene set. ITS sizes range from 1 to 9.68 and CTS sizes range from 1 to 37. In eight out nine data sets, the mean ITS sizes are below 4, but the instances in the brain cancer data set GSE84465 are more difficult and have much larger optimal solutions. Target genes or target gene families that are shared across datasets of the same cancer type include *TNR* and across different cancer types include *KRT16* and the *SOX* family and the *MAGE* family.

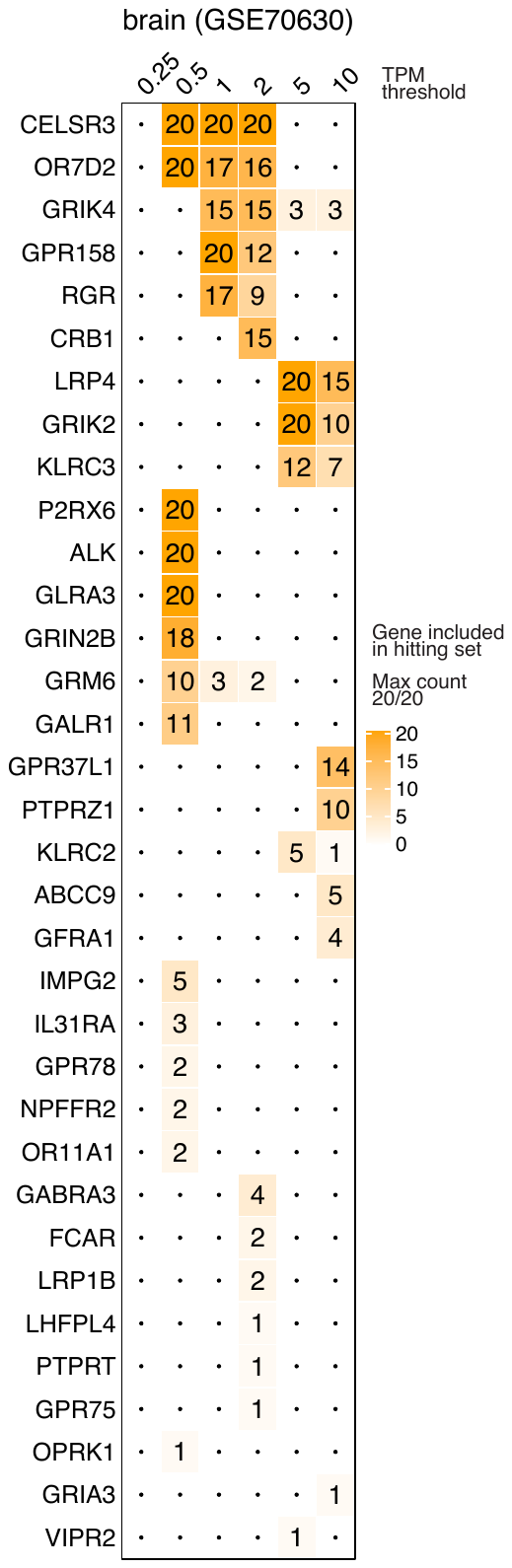

Figure S26. Frequency of genes found in the global hitting set of brain dataset (GSE70630).
The heatmaps represent the number of times a gene (cell-surface receptor) was observed in the optimal cohort target set (across 20 replicates) when the input target genes were sampled based on different thresholds of their mean expression levels across multiple healthy tissues (see Methods).

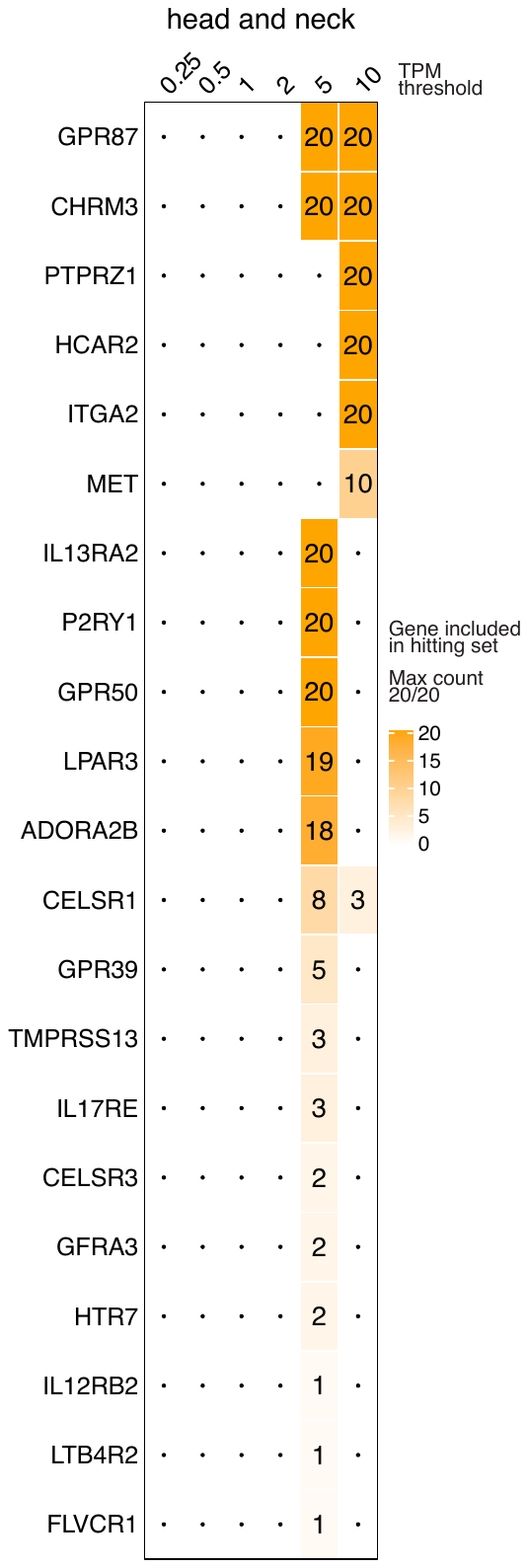

Figure S27. Frequency of genes found in the optimal cohort target set of head and neck dataset. The heatmaps represents the number of times a gene (cell-surface receptor) was observed in the optimal cohort target set (across 20 replicates) when the input target genes were sampled based on different thresholds of their mean expression levels across multiple healthy tissues (see Methods).

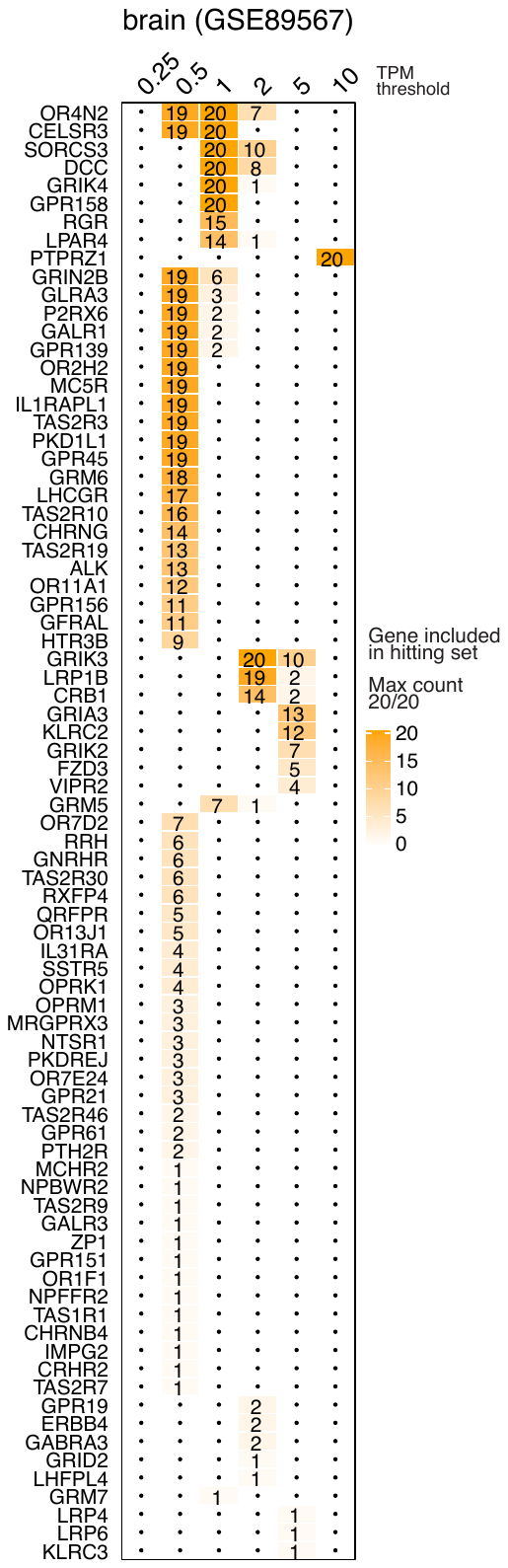

Figure S28. Frequency of genes found in the optimal cohort target set of brain cancer dataset (GSE89567). The heatmaps represents the number of times a gene (cell-surface receptor) was observed in the global hitting set (across 20 replicates) when the input target genes were sampled based on different thresholds of their mean expression levels across multiple healthy tissues (Methods).

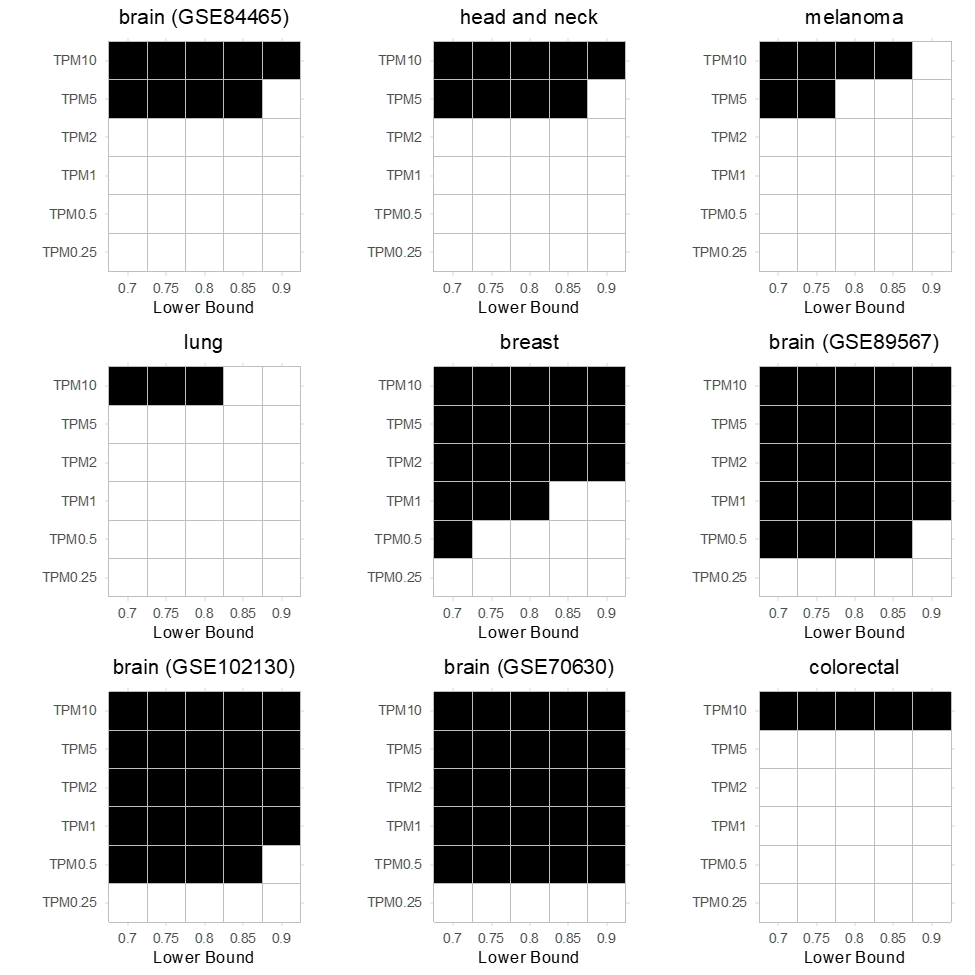

Figure S29. Availability of feasible solutions for target genes obtained through different cross tissue expression thresholds and lower bound parameter settings. Each heatmap represents a different dataset, the title of the heatmap represents the dataset/cancer type used. If a cancer type has more than one dataset representing it than the dataset identifier is provided in brackets. The x-axis denotes different lower bound parameter values and the y-axis denotes different thresholds used to identify target genes with low expression across tissues. Black color indicates the presence and white absence of a feasible solution. The results are based on one (random) out of 20 replicates for each dataset.

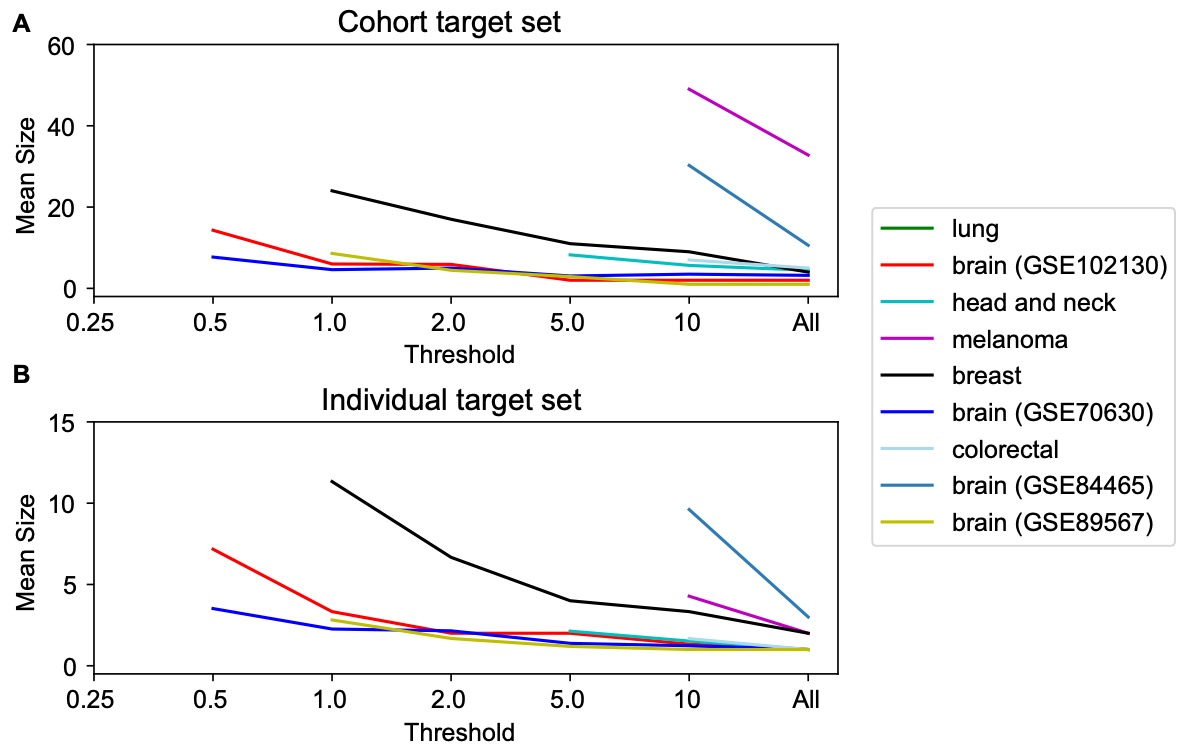

Figure S30. The sizes of the optimal cohort and target sets decrease as the size of the set of available gene targets increases. Each line in the plot represents the variation in the size of the optimal target set at cohort level (A) and individual patient level (B), when the number of target genes differ according to their expression levels (Table 2). The color of the line representing different cancer type is defined in the legend. The x-axis denotes the different thresholds used to identify target genes with low expression across tissues. The y-axis denotes the mean size of the individual target set among 20 replicates.

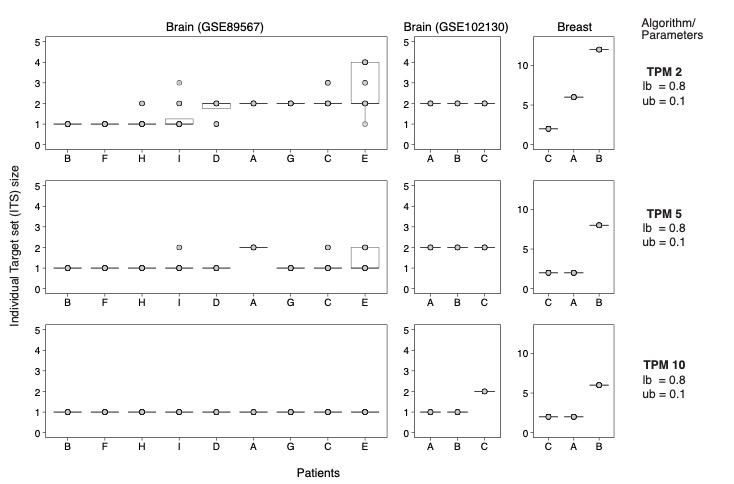

Figure S31. The distribution of ITS sizes across three different cancer types, at our baseline parameter setting and different cross tissue expression thresholds. The x-axis represents the individual patients within the dataset, y-axis represents the ITS size, and the boxplots represent the ITS sizes from 20 replicates.

Table S5: Summary of optimal solution sizes based on 533 potential CAR-T targets suggested by McKay et al.^39^. CTS is mean of 20 replicates. ITS is mean over all patients over 20 replicates. The rightmost column shows the genes that appear in an optimal CTS for at least 16/20 (80%) of replicates.

| GEO | Cancer Type | CTS | ITS | Genes found in ≥ 80% of the optimal CTS |
| --- | --- | --- | --- | --- |
| E-MTAB-6149 | Lung | 7.25 | 1.81 | *NPW, MAGEA4, SFTPB, SOX2, TUBB2B* |
| GSE102130 | Brain | 1 | 1 | *TNR* |
| GSE103322 | Head and neck | 6.2 | 1.3 | *MAGEA4, RIPK4, SOX2, COL22A1, CT45A5, KRT16* |
| GSE115978 | Melanoma | 37 | 3.55 | *TUBB2B, ZFAT, CLGN, IGF2BP1, LAMA1, MFAP2, MIA, RIPK4, MAGEA4, GAPDHS, CC2D2A, CCDC40, CTAG1B, SSX2B, TUBB4A, C2, BNC2, HSPA12A, MAGEC2, GDF15, SSX2, PEG10, MMP11, KALRN, PTGES, HSPA4L, FOSL1, LIN28A, CPB2, DAZ4, C16orf95, HERC5, SOX11, KRT16, AURKA, LCN10, FAM216A, TL10* |
| GSE118389 | Breast | 10 | 4.33 | *UBA6, PEG10, HLTF, KRT16, P2RY8, SLC26A4, DYRK4, ESR1, RND2, SOX11, CEACAM8, CC2D2A, MIA, PCDH11Y, CD79B, CCDC114, MCTP1* |
| GSE70630 | Brain | 2.7 | 1.01 | *ATCAY, TNR* |
| GSE81861 | Colorectal | 5 | 1.33 | *GDF15, C2, MCTP1, UBA6, HOOK1* |
| GSE84465 | Brain | 30.3 | 9.68 | *KCNN4, OTP, SOX2, C8orf48, KITLG, COL22A1, ANGPT1, SOX11, CFAP126, DCLK2, DNAH6, ESR2, SFRP4, MMP7, LEF1, HSD17B3, MUC12* |
| GSE89567 | Brain | 2 | 1.01 | *SOX11, TNR* |

**12. Further Analysis of Genes in Optimal Solutions for Different Data Sets and Parameter Setting not Covered in the Main Text**

By solving the same treatment optimization problem in multiple data sets from different tumor types and different labs, we can get a sense of which target proteins merit the most experimental effort to design nanoparticles or other treatments targeting that protein. Tables S6 through S8 show the genes appearing most often in the optimal solutions of the fair hitting set problem with $\alpha=0, 1, ..., 5$ using either 58 targets with known peptide-mimicking ligands (Tables 3 and 4) or 1269 cell-surface receptor targets (see Methods) and either all 9 data sets or only 4 brain cancer data sets that include both cancer and non-cancer cells. For each value of $\alpha$, we sampled 20 replicates as described in Methods. Thus, there are 1080 instances among the nine data sets or 480 instances among the four brain cancer data sets. For the 1269 targets, all 1080 instances were feasible, while when we restricted the universe to 58 targets, 990 out of 1080 instances were feasible.

Table S6. Genes occurring most in the highest proportion of optimal solution of instances of the fair hitting set problem for tumor cells and non-tumor cells, 58 targets, in the six datasets that are feasible at baseline settings and $\alpha=0, 1,..., 5 , ub=0.1, lb=0.8.$We used 20 replicates for each value of the parameters.

| Proportion of optima including gene | Gene Symbol |
| --- | --- |
| 0.661 | *EGFR* |
| 0.362 | *CDH1* |
| 0.334 | *FGFR2* |
| 0.319 | *MET* |
| 0.264 | *CDH2* |
| 0.223 | *EPHB1* |
| 0.218 | *ERBB2* |
| 0.201 | *VIPR2* |
| 0.177 | *APP* |
| 0.1704 | *CD93* |
| 0.1677 | *CXCR4* |
| 0.167 | *FGFR4* |
| 0.167 | *MUC1* |
| 0.150 | *ACVR2B* |
| 0.140 | *LRP1* |
| 0.139 | *PLAUR* |
| 0.113 | *PTPRJ* |
| 0.091 | *GRPR3* |
| 0.079 | *CD274* |
| 0.071 | *PROM1* |

Table S7. Genes occurring most often in the optimal solution of instances of the fair hitting set problem in nine cancer data sets for tumor cells and non-tumor cells, 1269 targets, $r=2, ub=0.1, lb=0.8$ and $\alpha=0, 1, ... , 5$. The maximum value possible in the left column is 1080 because there are 9 data sets, 6 values of $\alpha$ and 20 replicates for each data set and each value of $\alpha$.

| Number of Times | Gene Symbol |
| --- | --- |
| 388 | *PTPRZ1* |
| 225 | *CLDN4* |
| 165 | *CXADR* |
| 140 | *EPHB4* |
| 139 | *NTRK2* |
| 135 | *EGFR* |
| 134 | *SLC2A1* |
| 122 | *ERBB3* |
| 122 | *IL17RD* |
| 120 | *EDNRB* |
| 120 | *FRRS1* |
| 120 | *GPR137* |
| 120 | *INSR* |
| 120 | *LGR5* |
| 120 | *PLAUR* |
| 120 | *SORL1* |
| 118 | *TNFRSF14* |
| 116 | *EDA2R* |
| 116 | *GABRE* |
| 113 | *GPR87* |
| 100 | *GPRR* |
| 100 | *PTPRJ* |
| 91 | *CD44* |
| 90 | *CLDN3* |
| 82 | *CHRNA1* |
| 82 | *P2RX6* |
| 80 | *MCAM* |
| 77 | *FGFR1* |

Interestingly, at least three of the ten most frequently occurring targets among the 1269 targets (Table S7) already have known mimicking peptides: *EPHB4*, *EGFR*, and *ERBB2* (Tables 3 and 4) and another five with known mimicking peptides, *PLAUR*, *PTPRJ, CD44, MCAM, FGFR1*, rank between 14th and 30th.

Table S8. Genes occurring most often in the optimal solution of instances of the fair hitting set problem in four brain cancer cells for tumor cells and non-tumor cells, 1269 targets, and $\alpha=0, 1,\ldots, 5.$ The maximum value possible in the left column is 480.

| Number of Times | Gene Symbol |
| --- | --- |
| 303 | *PTPRZ1* |
| 135 | *EGFR* |
| 90 | *CD44* |
| 57 | *EPHB4* |
| 53 | *PILRB* |
| 47 | *LRP4* |
| 39 | *FZD7* |
| 35 | *OSMR* |
| 35 | *RAMP3* |
| 32 | *NTRK2* |
| 30 | *F2R* |
| 27 | *MET* |
| 24 | *GRIK2* |
| 22 | *CD163L1* |
| 20 | *SMO* |
| 17 | *GFRA1* |
| 15 | *HRH1* |
| 13 | *CELSR3* |
| 13 | *DDR2* |
| 13 | *ITGA2* |

The analysis in Tables S6-S8 look at the most common genes in isolation. One way to look at combinations of genes is co-occurrence networks as shown in main Figures 6C-6F. Another way is to use existing knowledge about protein functions and protein-protein interactions to examine whether the genes selection in optimal target sets are functionally connected. In the co-occurrence networks, the genes are deliberately and visually weighted by frequency of occurrence. In contrast, for the functional analyses, we use an unweighted list of the 25 or 30 most frequent genes. We did the functional analysis can be done via the STRING database and its many associated analysis tools for pathway enrichment. As visual examples of STRING analyses, we show in Figures S32-S35 protein-protein interaction networks for the (genes most commonly in optimal solutions) for the same four data sets used in main Figures 6C-6F. The tightly connected network of 15/25 genes selected for the head and neck cancer data set is particularly striking (Figure S33).

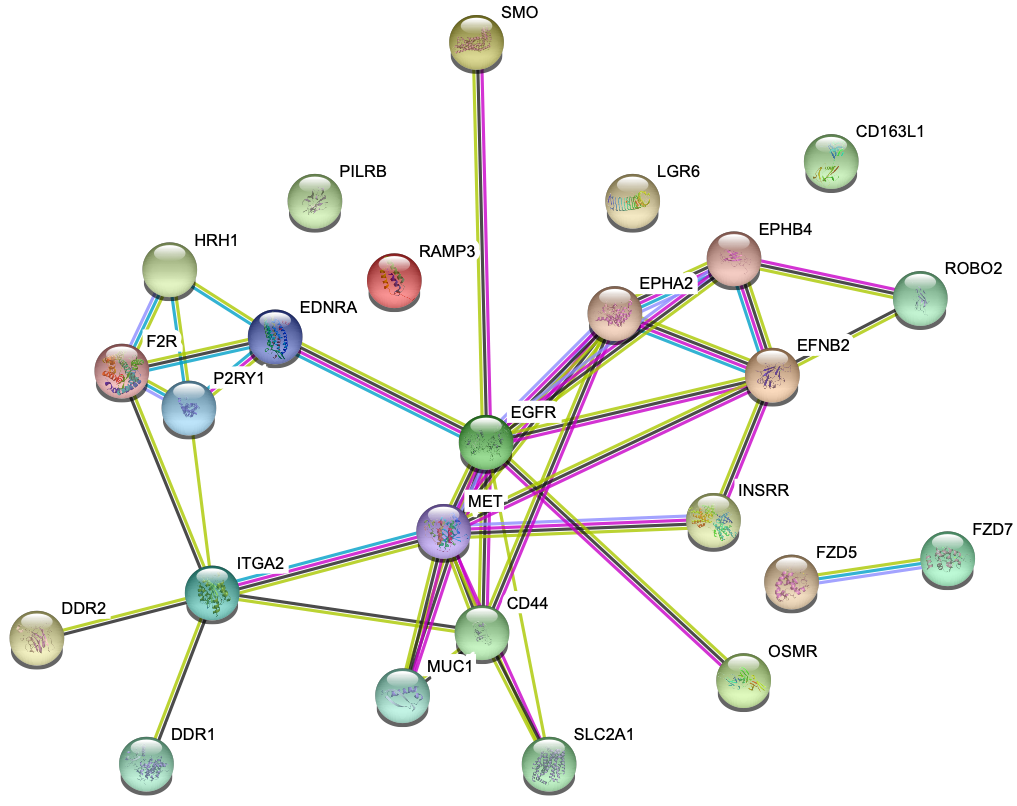

Figure S32. Protein-protein interaction network inferred and drawn by STRING^40^ for the 25 genes most commonly occurring in optimal cohort target sets for the brain cancer (GSE84465) dataset featured in the main document, as a weighted average over 50 optimal solution for each of 20 replicates at baseline settings.

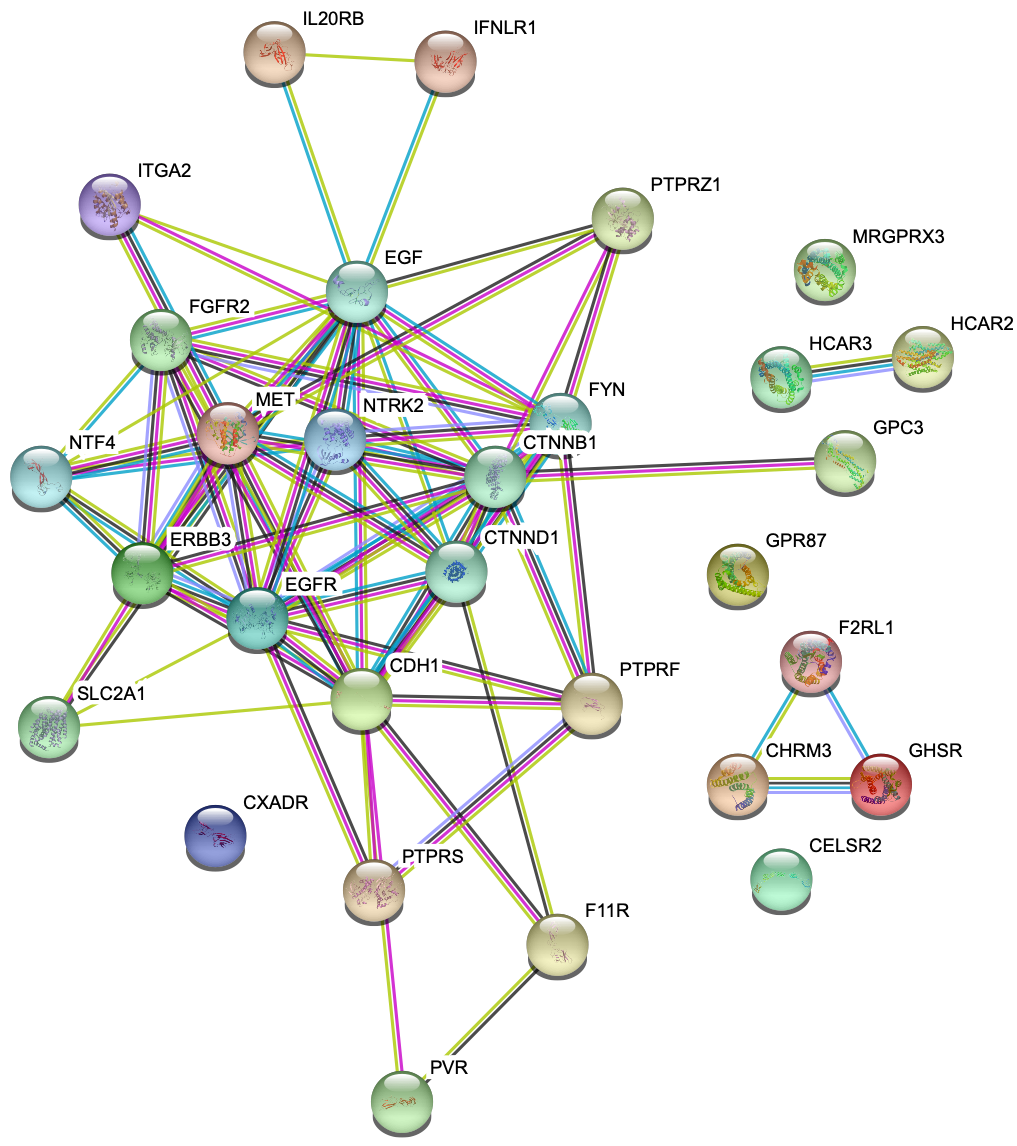

Figure S33. Protein-protein interaction network inferred and drawn by STRING^40^ for the 25 genes most commonly occurring in optimal cohort target sets for head and neck cancer dataset, as a weighted average over 50 optimal solution for each of 20 replicates at baseline settings.

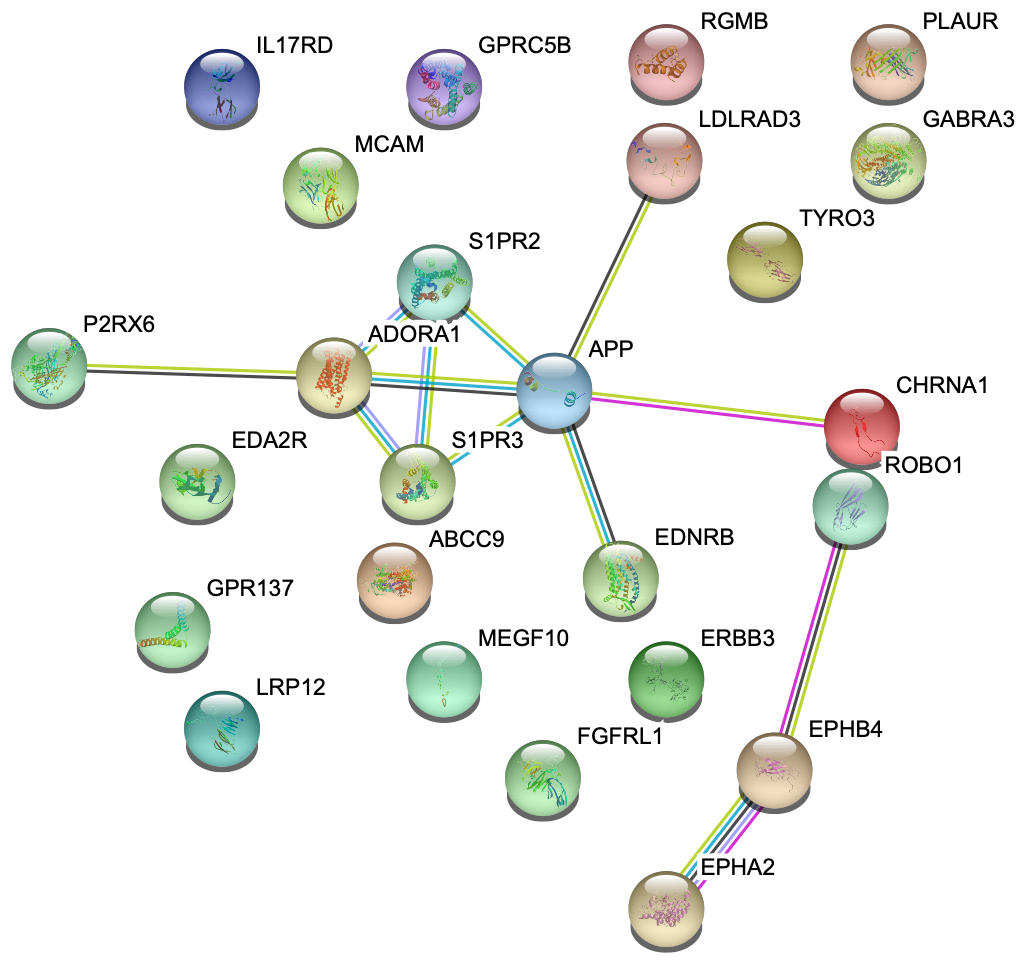

Figure S34. Protein-protein interaction network inferred and drawn by STRING^40^ for the 25 genes most commonly occurring in optimal cohort target sets for melanoma dataset, as a weighted average over 50 optimal solution for each of 20 replicates at baseline settings.

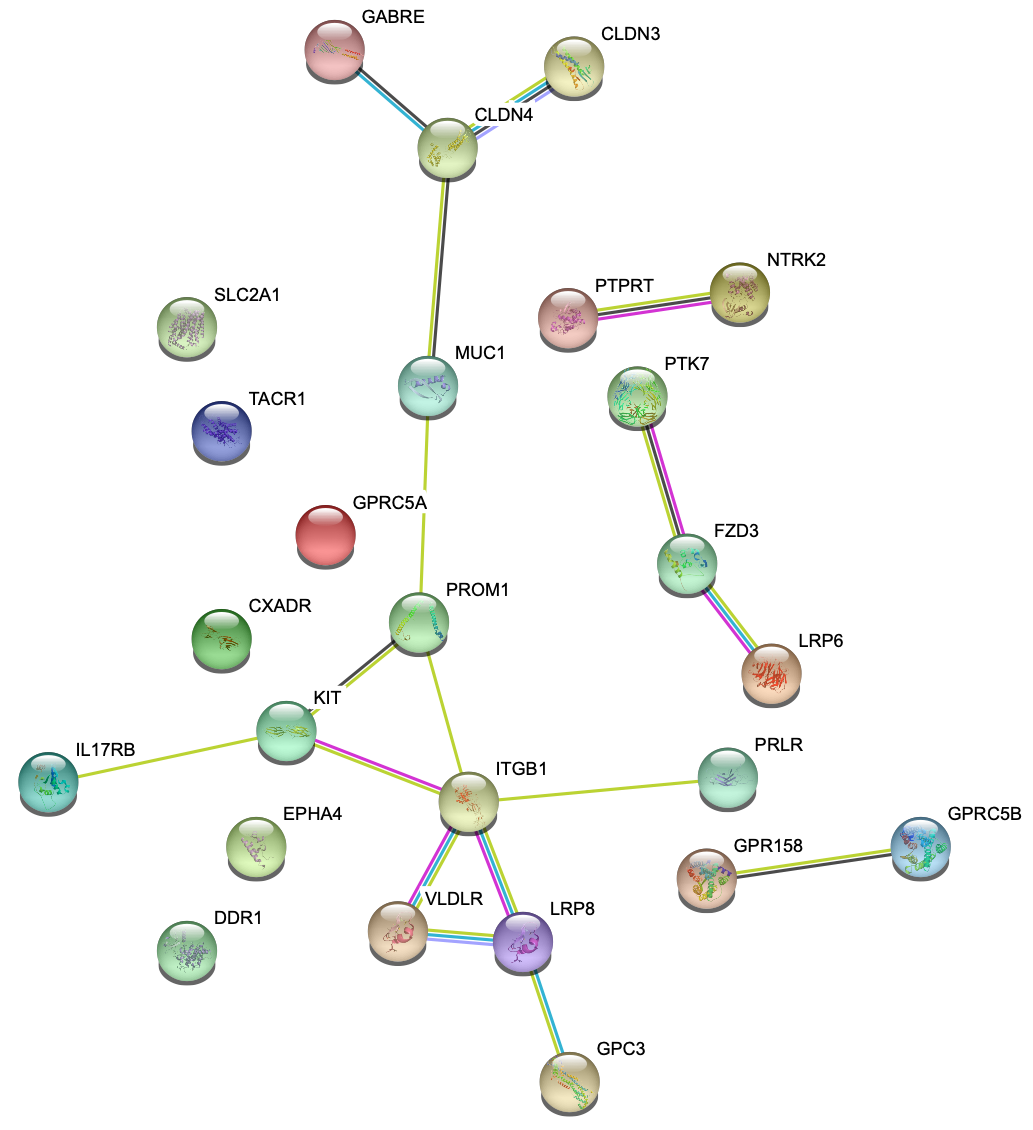

Figure S35. Protein-protein interaction network inferred and drawn by STRING^40^ for the 25 genes most commonly occurring in optimal cohort target sets for lung cancer dataset, as a weighted average over 50 optimal solution for each of 20 replicates at baseline settings.
